# BaalChIP: Bayesian analysis of allele-specific transcription factor binding in cancer genomes

**DOI:** 10.1101/093393

**Authors:** Ines de Santiago, Wei Liu, Martin O’Reilly, Ke Yuan, Chandra Sekhar Reddy Chilamakuri, Bruce A.J. Ponder, Kerstin B. Meyer, Florian Markowetz

## Abstract

Allele-specific measurements of transcription factor binding from ChIP-seq data are key to dissecting the allelic effects of non-coding variants and their contribution to phenotypic diversity. However, most methods to detect allelic imbalance assume diploid genomes. This assumption severely limits their applicability to cancer samples with frequent DNA copy number changes. Here we present a Bayesian statistical approach called BaalChIP to correct for the effect of background allele frequency on the observed ChIP-seq read counts. BaalChIP allows the joint analysis of multiple ChIP-seq samples across a single variant and outperforms competing approaches in simulations. Using 548 ENCODE ChIP-seq and 6 targeted FAIRE-seq samples we show that BaalChIP effectively corrects allele-specific analysis for copy number variation and increases the power to detect putative cis-acting regulatory variants in cancer genomes.

## Background

Allele-specific measurements of transcription factor binding from ChIP-seq data have provided important insights into the allelic effects of non-coding variants and their contribution to phenotypic diversity [1, 2, 3, 4, 5]. Since the majority of disease risk-associated SNPs occur in non-coding DNA, many of them might disrupt cisregulatory elements. Thus, a direct method to identify functional SNPs is to use the information obtained from allelic-specific binding (ASB) of transcription factors.ChIP-seq data are commonly used to study allele-specific binding of proteins at heterozygous non-coding SNPs and to infer putative effects of such variants on gene regulation. Allele-specific mapping corrects for environmental sources of variation that alter gene regulation, since both alleles are assayed in the same cellular context.

### Existing approaches to infer allelic imbalance

Previous studies have used ChIP-seq and RNA-seq to identify ASB and allele-specific expression (ASE). These studies have described methods to address technical and methodological biases such as the sequence context of a SNP [6], alignment biases to the reference allele [7, 8], and the issue of increasing detection power by combining multiple SNPs in the same gene [9] or across multiple ChIP-seq samples [10]. However, all of these approaches are designed to examine the allelic imbalances in diploid samples and do not address copy number differences between the two alleles, an ubiquitous feature of cancer genomes (Figure S1).

Few papers in the literature have tried to address confounding effects arising from copy number changes. Some studies have analyzed tumor and normal samples without making any adjustments to the applied methodology, which risks identifying false positives where the detected ASE is mainly due to copy-number alterations [11]. Others have removed all sites overlapping copy number variants [7, 12, 13], or used the genomic DNA allelic ratios to correct for the observed allelic imbalances [14, 15] or to remove SNPs with allelic imbalances detected in the control input DNA [2] which is only feasible when the coverage at any assayed heterozygous site is high (> 20x if a binomial test is to be applied to reach adequate statistical power [9, 12]). Recently, Liu *et al.* [16] developed cisASE, a likelihood-based method for detecting allele-specific expression. cisASE has been shown to successfully correct for copy number alterations that hinder the identification of ASE in RNA-seq data. Table S1 presents a summary of the different strategies of allelic-specific mapping analyses to deal with regions of altered copy-number.While the analysis of allele-specific expression and binding are conceptually similar, in practice methods tailored to either ASE or ASB are not easily exchangeable, because the data types and thus the biological and statistical assumptions are different.

### Bayesian analysis of allele specific binding

To address the wide range of biases that can affect the detection of ASB from ChIP-seq data we developed BaalChIP (Bayesian Analysis of Allelic imbalance from ChIP-seq data) (Figure 1). BaalChIP makes it possible to quantify the significance of the allelic imbalance from ChIP-seq data in cell lines with genomic aberrations, which is a major advance over previous approaches. BaalChIP combines several important features: First, we use several strategies to rigorously account for allelic mapping bias, including filtering of SNPs in problematic alignment regions as well as simulations to identify reads with increased risk of mapping bias. Second, we implement a beta-binomial model for allelic read counts. This model accounts for the fact that the observed variance in the data is larger than expected from a standard binomial model, a phenomenon known as overdispersion [9, 17, 18]. Third, we take advantage of the fact that multiple transcription factor ChIP-seq data may be available for the same SNP to improve ASB detection. Finally, we use a Bayesian framework to account for the influence of the background allele composition and the reference mapping bias on the observed ChIP-seq allelic read count.

**Figure 1.**
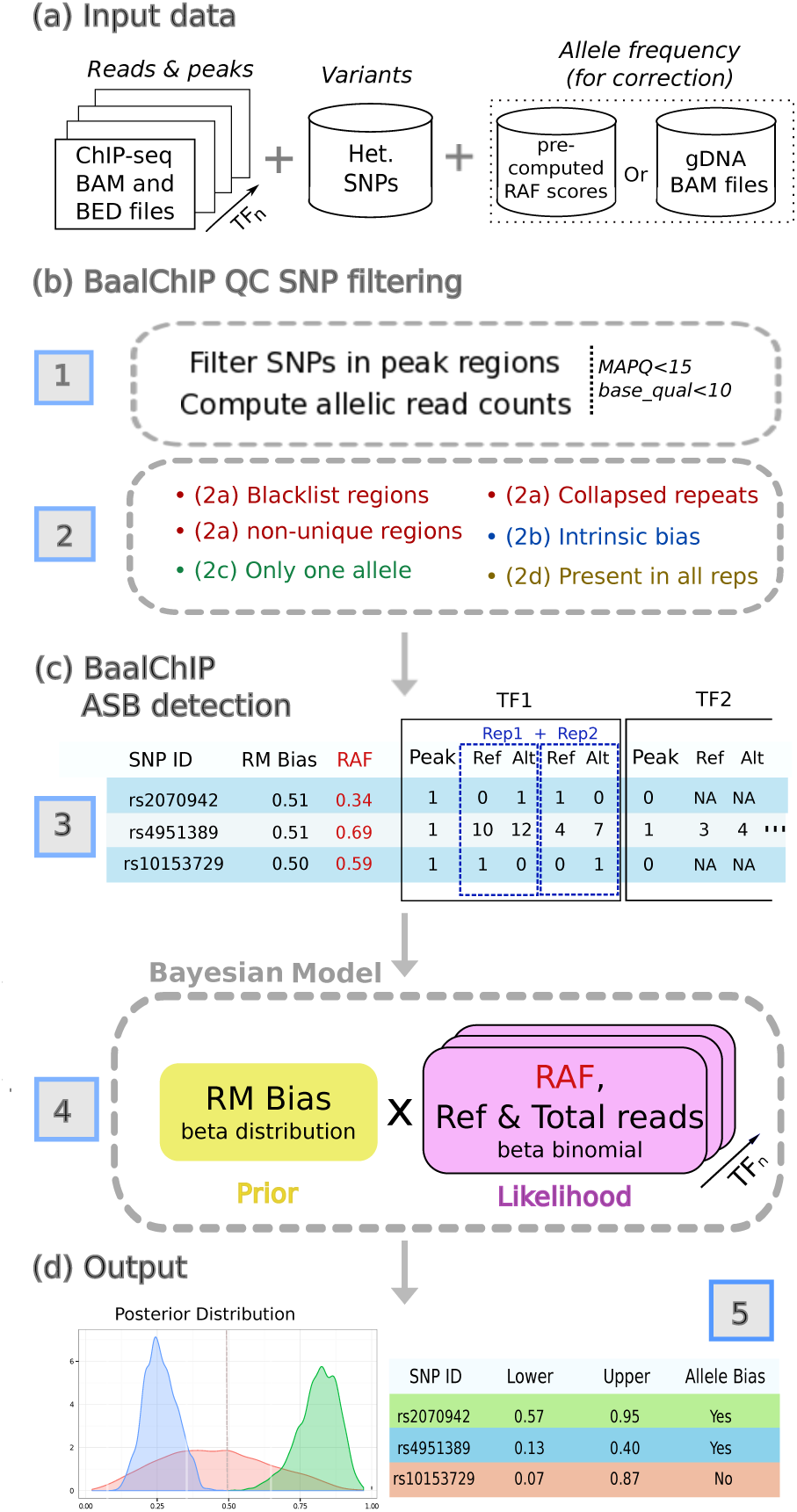
Description of BaalChIP model frame work. (a) The basic inputs for Baal are the ChIP-seq raw read counts in a standard BAM alignment format, a BED file with the genomic regions of interest (such as ChIP-seq peaks) and a set of heterozygous SNPs in a tab-delimited text file. Optionally, genomic DNA (gDNA) BAM files can be specified for RAF computation, alternatively, the user can specify the pre-computed RAF scores for each variant. (b) The first module of BaalChIP consists of (1) computing allelic read counts for each heterozygous SNP in peak regions and (2) a round of filters to exclude heterozygous SNPs that are susceptible to generating artifactual allele-specific binding effects. (3) the Reference Mapping (RM) bias and the Reference Allele Frequency (RAF) are computed internally and the output consists of a data matrix where RM and RAF scores are included alongside the information of allele counts for each heterozygous SNP. The column ‘Peak’ contains binary data used to state the called peaks. (c) The second module of BaalChIP consists of calling ASB binding events. (4) BaalChIP uses a beta-binomial Bayesian model to consider RM and RAF bias when detecting the ASB events. (d) The output from BaalChIP is a posterior distribution for each SNP. A threshold to identify SNPs with allelic bias is specified by the user (default value is a 95% interval). (5) The output of BaalChIP is a credible interval (’Lower’ and ‘Upper’) calculated based on the posterior distribution. This interval corresponds to the true allelic ratio in read counts (i.e. after correcting for RM and RAF biases). An ASB event is called if the lower and upper limits of the interval are outside the 0.4-0.6 interval.

We applied BaalChIP to 548 ENCODE samples [19] obtained from a panel of 14 cell lines, representing different tissues and karyotypes, including 8 cancer cell lines (HeLa-S3, A549, MCF-7, T-47D, K562, HepG2, SK-N-SH, HL-60) and 6 non-cancer cell lines (H1-hESC, GM12878, GM12891, GM12892, MCF10A-Er-Src, IMR90), as well as 6 FAIRE-seq samples obtained from breast cancer cell lines (MDA-MB-134 and T-47D). We demonstrate that copy-number changes can easily give rise to spurious signals of allelic imbalance. We are able to integrate the quantitative information obtained from ChIP-seq data along with information of the background allele composition to accurately detect allele-specific binding and correct for artifacts caused by allele-specific losses or gains at the structural genomic level. Because of this integrated modeling, we were able to detect a large number of ASB events from ChIP-seq data generating a resource for future functional studies.

BaalChIP is implemented as an R [20] package and freely available from Bioconductor repository https://bioconductor.org/packages/release/bioc/html/BaalChIP.html [21].

## Results

### Overview of BaalChIP workflow

In this study we aim to devise a method that allows to correct for copy number changes and other biases in the analysis of ASB from ChIP-seq or similar data. The BaalChIP workflow requires three different sets of input data (Figure 1): the SNP variants file, ChIP-seq datasets (e.g. sets of BAM and BED files of ChIP-seq data obtained for different proteins that may or may not be grouped into replicates), and a corresponding set of genomic DNA files for each individual sample. The SNP variants file and the BED files are used to select the sites for the analysis, the ChIP-seq BAM files are used to compute the allelic read counts and the gDNA files will allow to determine reference-allele frequency (RAF) values to correct for effects of the background allele composition. Alternatively, RAF values may be directly precomputed from BAF (B allele frequency) scores (Figure 1a). It has been shown that removing duplicate reads can reduce technical sources of biases at ASB sites [22] and we suggest that BAM files are pre-processed to contain only uniquely aligned reads or flagged duplicated reads.

#### BaalChIP Quality Control

The BaalChIP workflow starts by first computing the allelic read counts for each assayed heterozygous SNP overlapping genomic intervals of interest (e.g. ChIP-seq peaks). By default, only uniquely mapped reads (MAQ > 15), and sites with base quality > 10 are used (Figure 1b). In the next step several quality control (QC) filters are applied to consider technical biases that may contribute to the false identification of regulatory SNPs (Figure 1b).

In a first round, BaalChIP excludes SNPs mapping to regions of known problematic read alignment: blacklisted regions [23], non-unique regions with genomic mappability score of less than 1 (based on the UCSC mappability track, 50bp segments) [22], and collapsed repeat regions [24]. The excluded regions contain genomic intervals that are frequently enriched for repetitive elements and frequently cause ambiguous read mapping and sequencing artifacts [25].

The second QC filter performs read-alignment simulations to consider an additional type of read mapping bias named the intrinsic bias [6, 17]. This bias occurs due to intrinsic characteristics of the genome that translate into different probabilities of read mapping. Even when reads differ only in one location, reads carrying one of the alleles may have a higher chance of matching multiple locations (i.e. have many repeats in the genome) and may therefore be mapped to an incorrect locus. This, in turn, results in the underestimation of read counts and may cause both false-positive and false-negative inferences of ASB.

The third QC filter selects those SNPs that pass all filters in all replicated samples, provided that replicated samples exist. This step will mainly remove SNPs in regions where the ChIP-seq signal is not consistently detected across all replicates (for instance when coverage is zero in one of the replicates).

The final QC filter consists of removing possible homozygous SNPs by removing any site where only one allele is observed [4, 26].

#### BaalChIP detection of allele-specific binding (ASB) events

The filtered SNPs and their allelic read counts are merged into a table with the total number of read counts in the reference (Ref) and alternative (Alt) alleles. No data is entered (missing data, NA) if a SNP did not pass the previously applied QC step for that sample (Figure 1c).

BaalChIP considers two additional biases that may lead to inaccurate estimates of ASB: the reference mapping (RM) and the reference allele frequency (RAF) biases. The RM bias occurs because the reference genome only represents a single “reference” allele at each SNP position. Reads that carry the “non-reference allele” have an extra mismatch to the reference sequence. Previous work has shown that this creates a marked bias in favor of the alignment of reads that contain the reference genome and could therefore affect the accuracy of allele-specific binding estimates [6].

The RAF bias occurs due to alterations in the background abundance of each allele (e.g. in regions of copy-number alterations) and the correction for this bias is one of the key features of BaalChIP. RAF values at each heterozygous variant are used in the model likelihood to correct of the observed ChIP-seq read counts relative to the amount of the reference allele. These are given as relative measures from 0 to 1, where values between 0.5 and 1 denote an underlying bias to the reference allele, and a value between 0 and 0.5 to the alternative allele. RAF scores can be obtained from SNP array B-allele frequencies (BAF) by assigning the correct reference and alternative alleles to allele A and B generic labels (*RAF* is equal to *BAF* if the reference allele corresponds to the B allele; *RAF* is equal to *1-BAF* if the reference allele corresponds to the A allele). Alternatively, if whole-genome DNA (gDNA) sequencing samples are given as an input, BaalChIP calculates the RAF values directly from the gDNA allelic read counts.

Finally, BaalChIP uses a beta-binomial distribution to model read count data therefore accounting for extra variability (over-dispersion) in allelic counts, a phenomenon that is often observed in sequencing data [9, 17]. The output of BaalChIP is a posterior distribution of the estimated allelic balance ratio in read counts observed after considering all sources of underlying biases (Figure 1c).

### Evaluation of BaalChIP performance

In a controlled simulation study, we compare BaalChIP with the two major competing methods to infer allelic imbalance: binomial test and iASeq [10]. The binomial test is the most frequently used method for the analysis of allele-specificity from ChIP-seq data [1, 2, 3, 4, 7]. In a recent study [15], biases caused by differences in copy number were accounted for by weighting the binomial null with the allelic ratios observed from the gDNA samples (the number of reads mapping to each allele in the genomic DNA). Therefore, to take these strategy into consideration, we set the null hypothesis on the probability of success to the estimated RAF bias, instead of 0.5. iASeq is another available method that has been shown to improve the detection of ASB [10]. The iASeq method uses a Bayesian framework to combine information from different experiments or replicates, and models the read counts with a beta-binomal distribution, instead of a simple binomial distribution, to account for extra-binomial variation caused by technical variability. Although iASeq was not originally designed to overcome copy-number biases, we included it in our simulation study because of its offered improvement of the traditional binomial test.

All three methods are tested on synthetic data sets with varying sequencing depth, number of TF binding to a SNP and copy number states. In addition, we also examine the robustness of the methods against a wide range of true allelic balance ratios. The allelic imbalance calling performance is assessed by ROC (Receiver operating characteristic) curves. In the absence of copy number changes, i.e. RAF is 0.5, the performances of the three methods are similar (Figure 2a, d and g).

**Figure 2.**
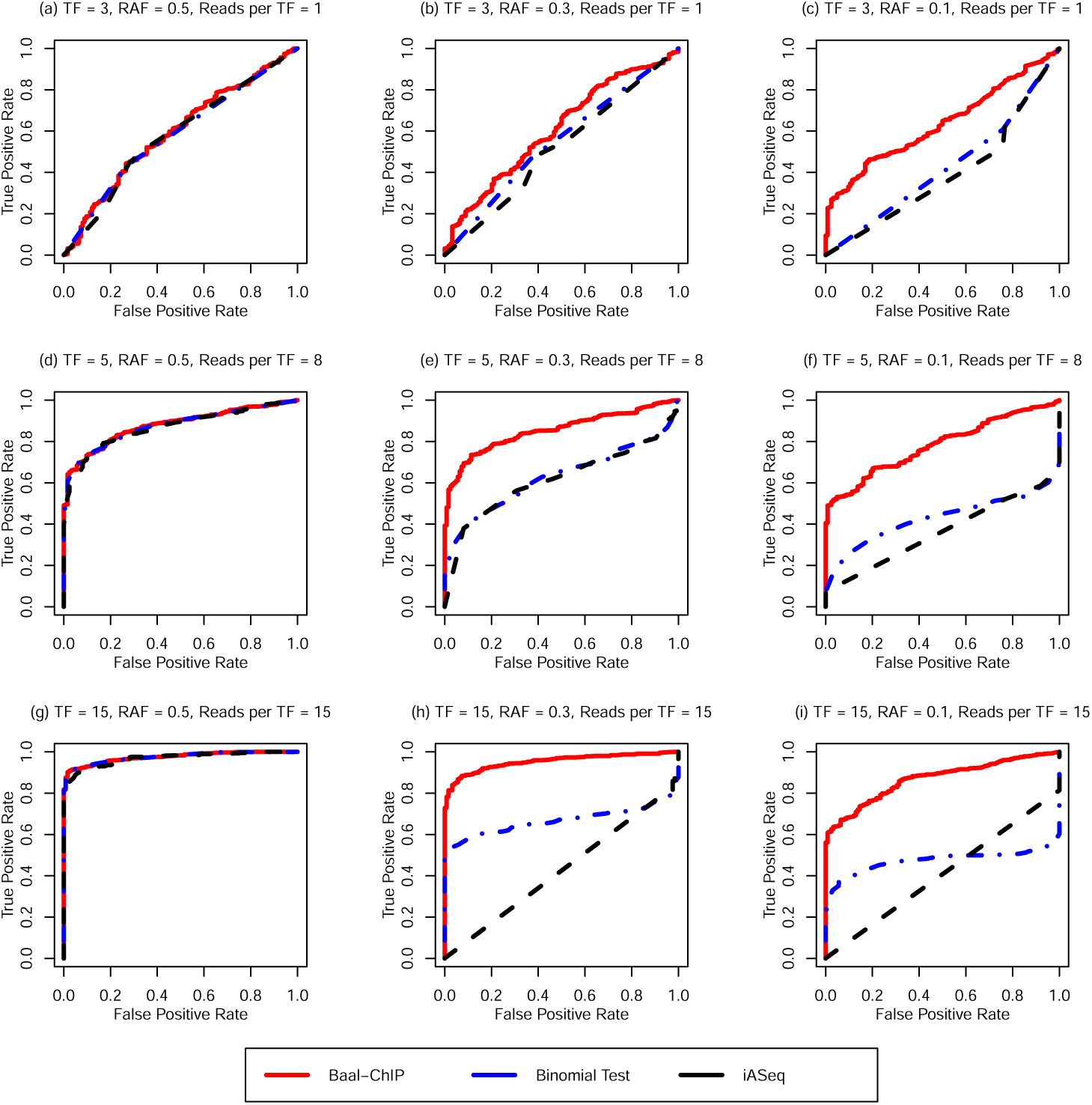
The ROC curve comparison between BaalChIP and other allele-specific SNP finding methods: binomial test and iASeq, using a simulated data set. The BaalChIP result is shown by solid red line. Binomial test and iASeq are shown in dashed blue and black lines. The number of TFs able to bind at a given SNP and reads per TF are increasing from 3 to 15 and 1 to 15, respectively. RAF is decreasing from 0.5 to 0.1.

In the presence of copy number changes, BaalChIP shows significant improvements in the identification of true ASB events. In Figure 2 we highlight results on data sets with RAF set to either 0.3 or 0.1, which represent modest and severe background allelic imbalance due to copy number changes. Significant improvements can be found across varying sequencing depth and across the number of TFs considered. Particularly, in the data sets with modest copy number changes, the performance of BaalChIP is similar to its performance in the absence of copy number changes, whereas the performance of both binomial test and iASseq suffers badly. When RAF is 0.1, BaalChIP is still able to achieve relatively good performance. In addition, the performance of BaalChIP benefits greatly from aggregating binding data for multiple TFs at the SNP (Figure S3).

Overall, the simulation results show that BaalChIP is robust and performs well across a wide range of variations in the data set. In particular, in regions with copy number changes BaalChIP shows significant improvements over the state-of-the-art baselines. Thus BaalChIP offers a robust analysis of ASB in samples obtained with frequent copy number changes.

### Case study 1: ENCODE data

We applied BaalChIP to 548 samples from the ENCODE project [19]. In total 271 ChIP-seq experiments were analyzed, assaying a total of 8 cancer and 6 non-cancer cell lines representing different tissues. The data contained either 2 or 3 replicates per experiment and 4 to 42 DNA-binding proteins per cell line (Table S3 and Figure S4 show a summary of the cell lines, tissues, and number of ChIP-seq experiments used in this case study).

The initial number of genotyped heterozygous SNPs per cell line ranged from 139K to 356K SNPs. We selected those SNPs that occurred within ChIP-seq peaks in the corresponding cell lines, which amounts to between 1.6% and 5.8% of all SNPs. To ensure a reliable set of heterozygous SNPs we applied the BaalChIP QC step with the default parameters and options. We removed an average of 30.4% of all SNPs that hit peaks (Figure S5b); 0.8% of the excluded sites were found in regions of problematic read alignments; 7.4% had biases in simulations (Figure S5a);16.4% were not consistent between replicates and 5.8% had only 1 observable allele (Figure S5c). After BaalChIP QC the number of heterozygous SNPs considered for downstream analysis ranged from 1,636-12,097 SNPs.

#### Allele-specific copy-number alterations change ChIP-seq read densities

First we examined of how allele-specific copy-number alterations affect the allelic ratios observed from ChIP-seq data (Figure 3). The relative presence of each allele is measured by the B allele frequency (BAF) obtained from SNP arrays [27].

**Figure 3.**
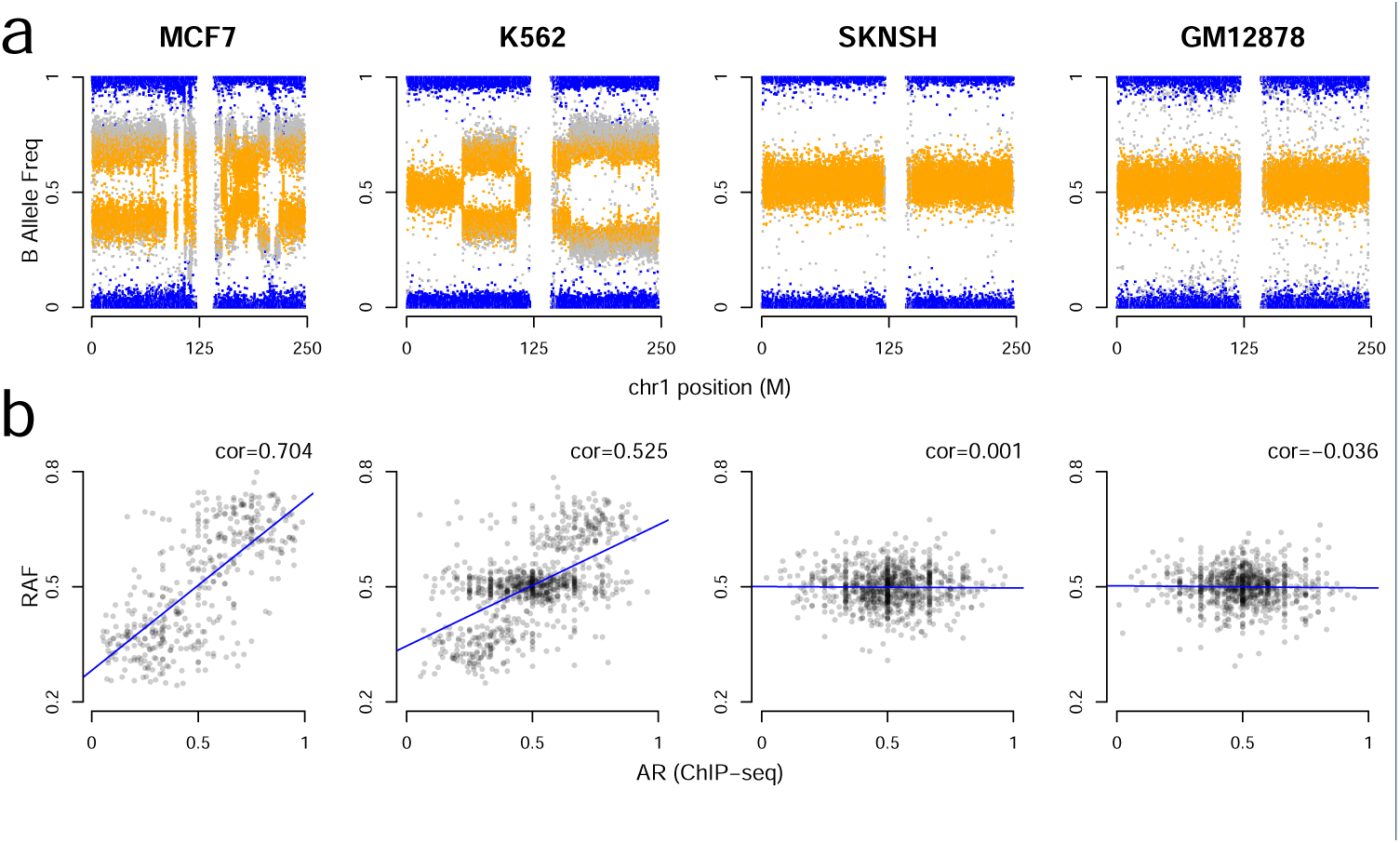
Example cancer and non-cancer cell lines SNP and ChIP-seq ENCODE data. (a) B allele frequencies (BAF) for chromosome 1 for three cancer cell lines (MCF-7, K562, SK-N-SH) and one non-cancer cell line (GM12878). Individual SNPs are colored according to genotype values: homozygous AA or BB (blue) and heterozygous AB (orange). (b) Correlations between the BAF values and the ChIP-seq allelic ratios (AR) of heterozygous SNPs. RAF corresponds to the BAF value with respect to the reference allele (RAF is equal to BAF if the reference allele corresponds to the B allele; RAF is equal to 1-BAF if the reference allele corresponds to the A allele). The fitted linear model (blue line) and the Spearman correlation coefficient (cor) show the relationship between BAF and ChIP-seq allelic ratios at heterozygous sites.

BAF values lie in the interval [0,1]. A BAF value of 0.5 indicates an equal presence of the two alleles and is the expected value for heterozygous sites (AB) in a diploid genome, while values of 0 and 1 indicate homozygous genotypes (AA and BB). In a normal, non-contaminated diploid sample, a BAF plot will have three bands, one centered at 0.5 for AB genotypes, and two bands at 0 and 1 for the AA and BB homozygous genotypes, respectively.

Figure 3a presents the BAF plots for chromosome 1 of three cancer cell lines (MCF-7, K-562 and SK-N-SH) and one non-cancer cell line obtained from normal blood (GM12878). As expected, the BAF plot of GM12878 is characterized by a typical diploid pattern of BAF values distributed around 0, 0.5 and 1 corresponding to the diploid genotypes AA, BB and AB. The few small deviations from these values can be attributed to germ line-based copy-number aberrations [28, 29]. The BAF plot for the SK-N-SH cell line, a glioblastoma cancer cell line known to have a near diploid karyotyope [30], is also relatively stable. In contrast, the MCF-7 and K-562 cancer cell lines show more complex BAF plots due to a variety of copy-number aberrations.

To evaluate the effects of allele-specific copy-number alterations on the ChIP-seq read densities, we compared for all four cell lines the BAF scores with ChIP-seq allelic ratios obtained at heterozygous SNPs (hetSNPs) (Figure 3b). We report ChIP-seq allelic ratios (AR) as the proportion of total reads carrying the reference allele. We observed a clear correlation between the BAF scores and the imbalance in the ChIP-seq allelic ratios in MCF-7 and K-562 cancer cell lines (Spearman *ρ* = 0.704 and 0.525, respectively), while this correlation is not observed in the near-diploid SK-N-SH (Spearman *ρ* = 0.001) and the normal GM12878 cell lines (Spearman *ρ* = 0.036).

The same effect can be observed in all other cell lines (Figure S6a) and is particularly striking in cancer cells with the highest proportion of extreme BAF scores (< 0.4 or > 0.6; Figure S6b). Taken together, these results demonstrate the need to adjust for the background genomic abundance of the alleles when attempting to identify putative cis-acting regulatory SNPs from ChIP-seq data. These data also support our assumption that for allelic changes that do not change TF affinity, the allele frequency in ChIP-seq data is proportional to the presence of these alleles in the genomic DNA.

#### Allele-specific amplification events explain most of the imbalances observed in ChIP-seq data in cancer cell lines

Having obtained a reliable set of heterozygous SNPs per cell line, we next applied the Bayesian framework implemented in BaalChIP to each of the 14 cell lines individually. To assess the importance of adjusting for copy-number biases, we performed the analysis with and without adjusting for the RAF bias. In a diploid sample, the allelic ratios (AR) are expected to be distributed around the 0.5 average, assuming that only a small proportion of sites carry an imbalance towards one of the alleles. However, as shown in Figure 3 copy-number alterations can affect read densities which will have an effect on ARs, in particular in cancer cells. Figure S7a shows the observed AR values in ChIP-seq data compared to the values before and after RAF correction for all analyzed cell lines. We observed that after RAF copy-number correction the ‘corrected allelic ratios’ become more evenly distributed around an average of 0.5. This effect is particularly notable in data obtained from cancer cell lines (Figure S7a).

Overall, we found 2,438 ASB sites across all cell lines (Table S4), with an average of 3.1% SNPs displaying allele-specific binding using our chosen cutoffs. Figure S7b demonstrates that the copy-number correction has a strong effect in cancer cell lines. In the most extreme cases the number of allele-specific imbalances is reduced by 4-fold. In normal cells, and in cancer cells with near diploid genomes, such as HL-60 and SK-N-SH [30, 31], the total number of identified ASB sites due to copy-number effects is much lower (Figure S7b). This is consistent with the fact that non-cancer cell lines carry far fewer copy number aberrations. Because ASB is more easily detected in regions of high sequence coverage due to increases in statistical power (Figure S8), we repeated the same analysis after selecting only SNPs with 30x-40x read coverage to control for read depth biases, and demonstrate that the same effect is verified (Figure S7c). Importantly, the data shown here demonstrates that adjusting for RAF is able to remove artifacts that are caused by copy number alterations in cancer cell lines.

We observed higher rates of ASB on chromosome X of female cell lines (GM12878, GM12892, IMR90, MCF10, MCF-7, SK-N-SH) than in autosomal chromosomes (Figure S9, *χ*^2^ test p-value < 2.2e-16 for chromosome X versus autosomal identified sites comparison for all 6 cell lines). These cases might be explained by the extent of X-inactivation. In normal tissue X-inactivation is random. However, in clonal cell lines the same X-chromosome will continue to be silenced and most X-linked genes are expressed in a mono-allelic fashion [32].

#### ASB events are consistent within and between cell lines

We evaluated the correlation of allelic ratios between pairs of biological replicates, between distinct proteins bound at the same site, and at identical ASB sites in different cell lines. The allelic ratios of all shared heterozygous SNPs are well correlated across biological replicates (Figure S10). Secondly, different proteins binding to the same SNP also display concordant allelic ratios (Figure S11) across cell lines. Our observations are consistent with the concordance of allele-specific binding of different co-bound TFs described in previous studies [1, 2, 3, 10, 12, 33, 34]. The Spearman correlation coefficients of all pairwise comparisons (between replicates and within cells between different DNA-binding proteins at the same SNP) are plotted as a boxplot in Figure S12a. Positive Spearman correlation coefficients are observed in every case, and in the majority of the cases we observe correlation coefficients > 0.8. Therefore, overall we find that BaalChIP generates highly reproducible results, across separate ChIP-seq experiments.

In addition, we analysed if heterozygous sites shared by cell lines had allelic ratios that were skewed towards the same allele or not. Of the identified ASB sites, only a small proportion (149 out of 2,438; 6.1%) were shared between cell lines (Figure S12b). Of these ASB sites, 91% (136 out of 149; Table S5) show an agreement in the direction of the allelic bias. Discordant cases may be explained by environmental or epigenetic factors, by different genomic context in different cells [1, 2], or by low sequencing coverage of ChIP-seq data. The high proportion of ASB with the same allelic bias further supports the robustness of BaalChIP.

#### Functional annotation of ASB sites

We then examined the genomic distribution of the identified ASB SNPs. A large proportion of ASB SNPs overlap introns and intergenic regions, and a considerable proportion is found at promoter-proximal regions, mainly reflecting the binding patterns of the ChIPed proteins (Figure S13). A large proportion of ASB SNPs overlap previously predicted enhancer regions and histone modifications associated with active enhancers (H3K4me1, H3K27ac), with an average of 70.2% of ASB SNPs occurring within cell-type specific putative enhancer regions (Figure S14). However, this enrichment is not significant when compared to non-ASB SNPs (Table S6, *χ*^2^ pvalue > 0.05), and it is possibly mainly reflecting the distribution of binding sites (ChIP-seq peak regions) from which the initial set of heterozygous SNPs was sampled.

Finally, to examine the putative functional mechanisms of ASB SNPs, we assessed if they modulated transcription factor binding affinity. We used predictions from HaploReg to assess if the observed ASB SNPs alter canonical transcription factor binding motifs. We found that 88% of ASB SNPs (range 83% to 92%) are motif disrupting SNPs but this enrichment was not statistically significant, since 85% of all tested SNPs under ChIP peaks were also found to alter motif scores. No significant difference in the magnitude of the change in binding affinity was observed between motif disrupting ASB SNPs when compared to all tested motif-disrupting SNPs, using the Kolmogorov–Smirnov test (Figure S15) To determine which TF-motifs are most likely to be disrupted, we grouped SNPs according to the DNA-binding proteins as identified by ChIP-seq peaks, and identified the top motifs disrupted by ASB SNPs when compared to non-ASB SNPs. In the majority of cases, the top disrupted motifs match the DNA-binding proteins used to generate the ChIP-seq data (Table S7). Overall, these results provide good evidence that the allele-specific binding we identify represents a true biological phenomenon.

### Case study 2: FAIRE-seq data

To demonstrate the generality of our approach, we applied BaalChIP to targeted FAIRE-sequencing data obtained from two breast-cancer cell lines, MDA-MB-134 and T-47D. FAIRE stands for Formaldehyde-Assisted Isolation of Regulatory Elements and is an effective method to identify DNA regions in the genome associated with open chromatin [35]. The method is based on the fact that the formaldehyde cross-linking is more efficient in nucleosome-bound DNA than it is in nucleosome-depleted regions of the genome. Thus, FAIRE-seq identifies regions of open chromatin. One advantage of FAIRE-seq over ChIP-seq is that the assayed chromatin is not limited to the location of specific DNA-associated proteins.

We chose to focus on the fraction of the genome that has been previously associated to breast-cancer risk. To do so, we selected 75 previously known breast-cancer risk regions (Table S8) [36] covering a total of 4.93Mb of the human genome. We performed targeted sequencing of 3 replicated FAIRE samples per cell-line and the correspondent genomic DNA (gDNA) controls. Targeted sequencing of the gDNA samples allowed us to determine with confidence the genotype of a high number of sites at the assayed breast cancer risk regions. We identified a total of 3208 and 1624 heterozygous SNPs in MDA-MB-134 and T-47D cells, respectively. In this dataset, the sequenced gDNA samples were used for the RAF correction step, i.e. allelic ratios at each SNP position were calculated directly from gDNA samples and used for bias correction. We first applied the BaalChIP QC pipeline to eliminate biases. We noticed that none of the SNPs in the selected targeted regions overlapped regions of potential problematic alignments, and only a small proportion of sites were eliminated during the BaalChIP QC step (< 0.3 %).

We observed high correlation between allelic ratios in the gDNA and FAIRE samples (Figure 4a), indicating that observed allele-specificity in FAIRE-seq samples is primarily due to copy-number alterations and must therefore be corrected for. Figure 4b shows the observed allelic ratios (AR) obtained in the FAIRE-seq samples compared to the values after correcting for gDNA. After correction the ARs become more evenly distributed around an average of 0.5. We found that approximately 0.65% (MDA-MB-134) and 0.56% (T-47D) of the tested sites in the selected risk regions were allele-specific. These correspond to a total of 21 sites in MDA-MB-134 and 9 sites in T-47D cell lines (Table 1).

**Figure 4.**
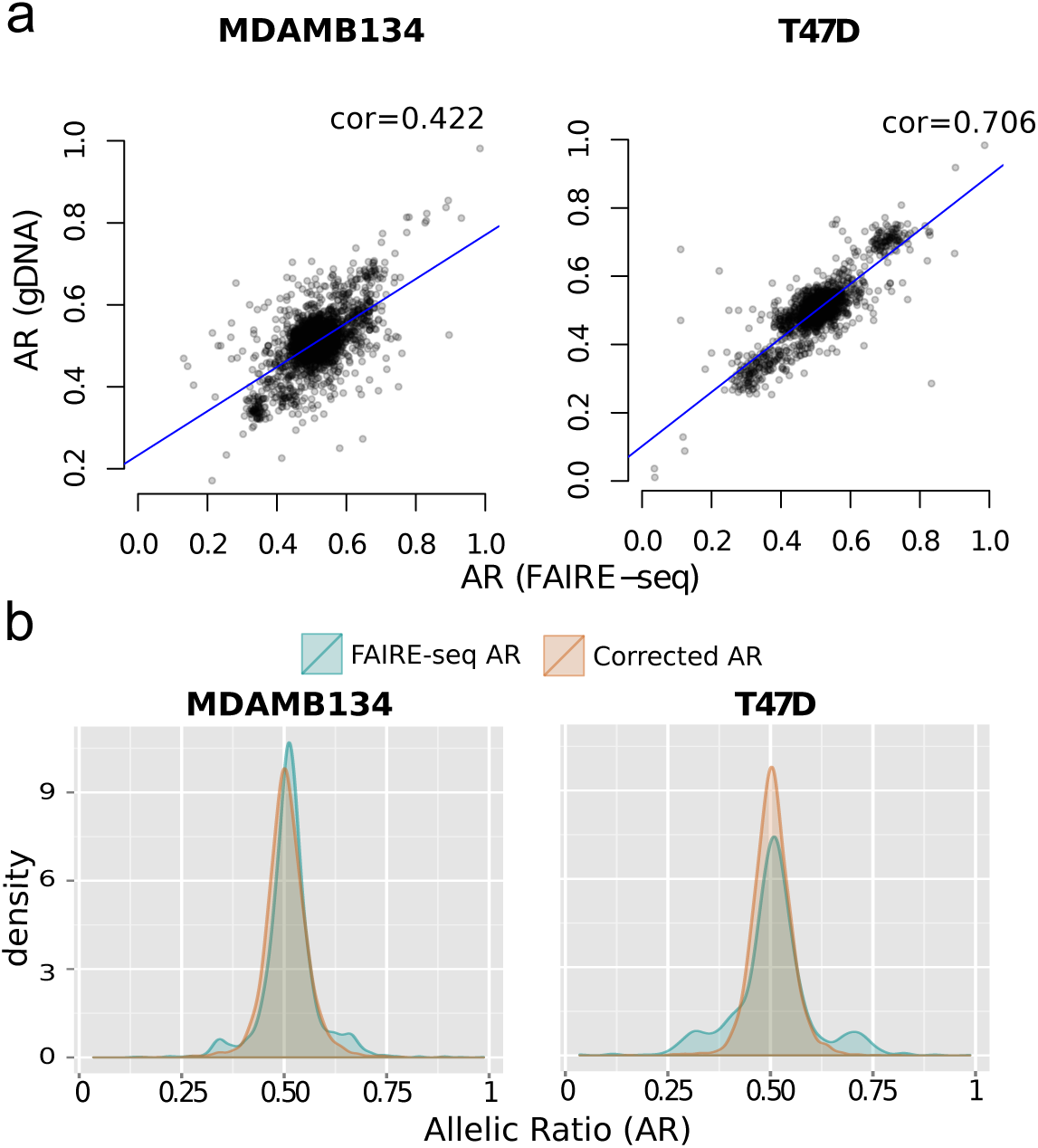
ASB detection from FAIRE targeted sequencing data. (a) Correlations between the allelic ratios obtained from genomic DNA and FAIRE-seq data. (b) Density plots showing the distribution of allelic ratios before (green) and after (orange) BaalChIP correction. The adjusted AR values were estimated by BaalChIP model after taking into account the RAF scores computed directly from the control genomic DNA samples.

**Table 1.** Heterozygous SNPs identified as allele-specific and the correspondent cell-lines and alleles in the FAIRE-seq dataset. CHROM, POS correspond to the coordinates in the hg19 human reference genome.

| ID | CHROM | POS | REF | ALT | ASB allele | Allelic Imbalance | GeneName | Location |
| --- | --- | --- | --- | --- | --- | --- | --- | --- |
| rs2373062 | 2 | 218322137 | G | C | T47D-G | 0.76 | DIRC3 | intron |
| rs9682063 | 3 | 4737842 | A | G | T47D-G | 0.34 | ITPR1 | intron |
| rs9292122 | 5 | 56087910 | G | A | T47D-G | 0.81 |  | intergenic |
| rs201314815 | 7 | 144115683 | C | T | T47D-T | 0.28 |  | intergenic |
| rs201949580 | 7 | 144115698 | T | G | T47D-G | 0.30 |  | intergenic |
| rs200312388 | 7 | 144115704 | T | G | T47D-G | 0.29 |  | intergenic |
| rs200496470 | 8 | 128323987 | A | G | T47D-G | 0.15 | CASC8 | intron |
| rs111929748 | 11 | 69379287 | G | T | T47D-T | 0.32 |  | intergenic |
| rs12939887 | 17 | 53162672 | A | G | T47D-A | 0.67 | STXBP4 | intron |
| rs72650670 | 1 | 114394681 | G | A | MDAMB134-G | 0.76 | PTPN22 | coding |
| rs116782319 | 5 | 158260122 | A | T | MDAMB134-T | 0.33 | EBF1 | intron |
| rs12670370 | 7 | 144118594 | C | A | MDAMB134-C | 0.79 |  | intergenic |
| rs2392922 | 8 | 129166840 | A | G | MDAMB134-A | 0.79 |  | intergenic |
| rs17793170 | 8 | 129205679 | G | A | MDAMB134-A | 0.33 |  | intergenic |
| rs2666763 | 10 | 22101608 | G | A | MDAMB134-A | 0.29 | DNAJC1 | intron |
| rs2941733 | 10 | 22241656 | T | C | MDAMB134-T | 0.74 | DNAJC1 | intron |
| rs4752570 | 10 | 123337066 | T | C | MDAMB134-C | 0.32 | FGFR2 | intron |
| rs2912780 | 10 | 123337117 | C | T | MDAMB134-C | 0.69 | FGFR2 | intron |
| rs2912779 | 10 | 123337182 | T | C | MDAMB134-T | 0.72 | FGFR2 | intron |
| rs2981579 | 10 | 123337335 | A | G | MDAMB134-A | 0.71 | FGFR2 | intron |
| rs2912778 | 10 | 123338654 | G | A | MDAMB134-G | 0.73 | FGFR2 | intron |
| rs11200017 | 10 | 123339685 | G | A | MDAMB134-A | 0.26 | FGFR2 | intron |
| rs2981578 | 10 | 123340311 | C | T | MDAMB134-C | 0.88 | FGFR2 | intron |
| rs11599804 | 10 | 123340664 | G | A | MDAMB134-A | 0.16 | FGFR2 | intron |
| rs34032268 | 10 | 123341525 | A | C | MDAMB134-C | 0.19 | FGFR2 | intron |
| rs2936870 | 10 | 123348902 | T | C | MDAMB134-T | 0.69 | FGFR2 | intron |
| rs3135718 | 10 | 123353869 | C | T | MDAMB134-C | 0.67 | FGFR2 | intron |
| rs112536831 | 19 | 17424329 | G | A | MDAMB134-G | 0.75 | DDA1 | intron |
| rs113207944 | 19 | 17438923 | C | T | MDAMB134-C | 0.74 | ANO8 | intron |
| rs10416361 | 19 | 44294617 | A | G | MDAMB134-A | 0.75 |  | intergenic |

Out of the 21 SNPs identified in the MDA-MB-134 cell line, 11 cluster in the 10q26.13 region. This cluster includes the rs2981579 SNP in the second intron of the FGFR2 gene (Table 1), which is the SNP with the strongest association with breast cancer in genome-wide analysis [37] In the 10q26.13 region, we identify two breast cancer risk SNPs, rs2981579 and rs2981578 with a strong allelic imbalance towards their risk alleles, rs2981579-A and rs2981578-C, respectively.

### Comparing BaalChIP to competing methods

We applied two competing methods, the binomial test and the iASeq [10] method, to the ENCODE and FAIRE-seq datasets.

When applying the iASeq method to the ENCODE dataset, the existence of missing data created an error. Missing data occur for those samples that do not contain ChIP-seq peaks at the SNP in question, or if the a SNP did not pass the previously applied QC step for that sample. To overcome this limitation of iASeq, we replaced every missing data point by zero, with the caveat that this *ad hoc* approach may create unknown biases in our results.

When applying the binomial test, for each cell line, we pooled read count data from different experiments and replicates. While this approach maximizes statistical power, it may mask heterogeneity in ASB obtained from different experiments. Previous studies have shown that at least 20x read coverage at a particular SNP position is necessary to reach adequate statistical power with the binomial test [9, 12]. Therefore, we restricted our analysis to SNPs with a coverage of at least 20 reads in the pooled data. We included two different ways of performing the binomial test, either by setting the probability of success to 0.5 or by weighting the binomial null with the RAF scores. In real data, unlike simulations, we are not able to access the true-positive and false-positive rates and compute ROC curves. For this reason, we mainly focused on testing whether BaalChIP is comparable to existing methods while improving the biased ASB detection in high copy-number regions (Figure 5a) or the problem of overdispersion in the data (Figure 5b).

**Figure 5.**
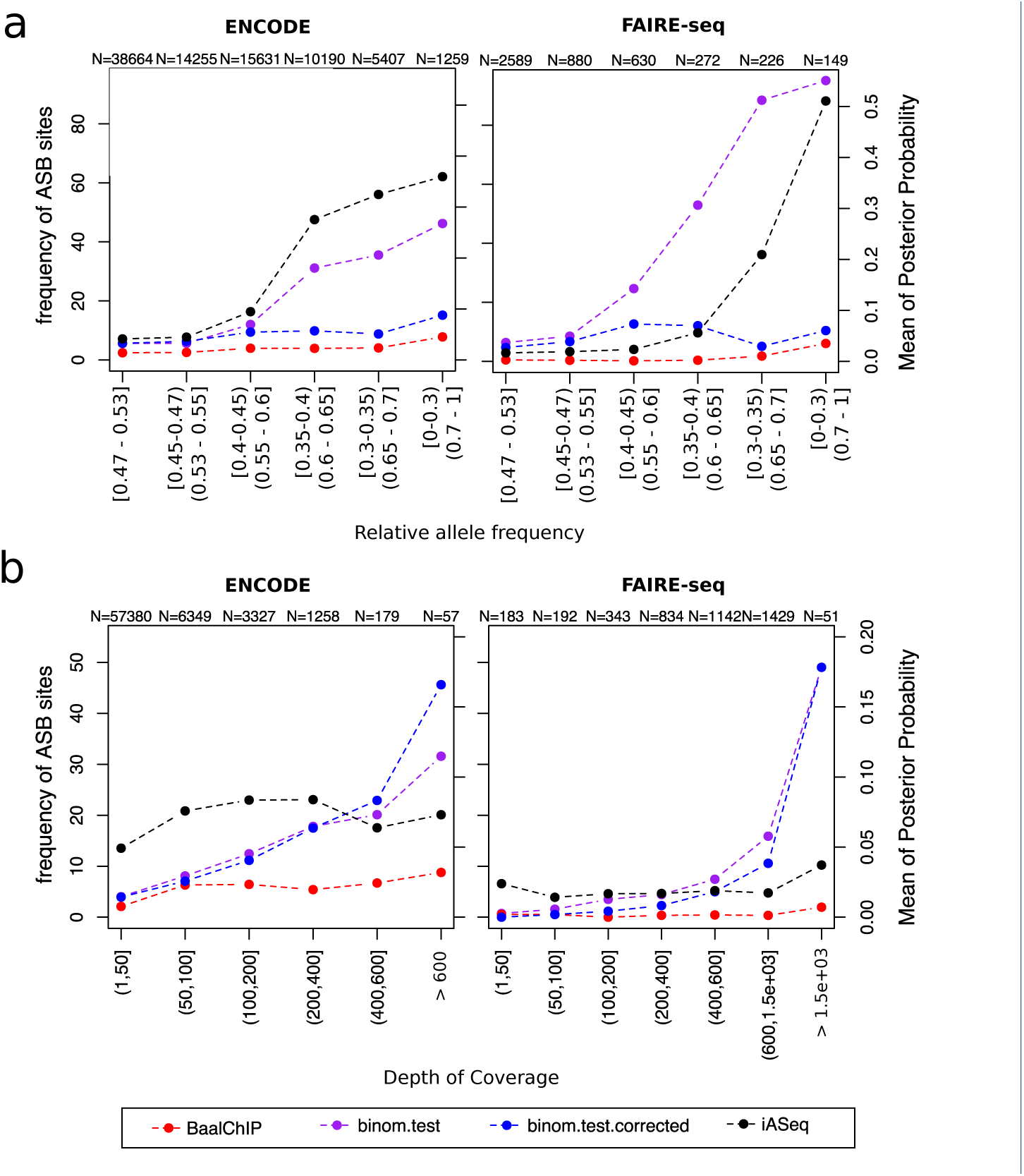
Comparison of BaalChIP with other available methods. Left y-axis corresponds to the frequency of ASB events called by BaalChIP or the binomial tests (red, blue and purple lines). Right y-axis corresponds to the mean of the posterial probability given by the iASeq method (black line). Small numbers at the top of the plot show the total number of tested heterozygous sites in each bin. (a) SNPs were grouped in bins of different RAF intervals. The RAF intervals are increasing in terms of distance to the diploid (RAF=0.5) value. The bionomial test (without RAF correction; purple line) and the iASeq methods (black line) are biased towards the detection of ASB events in regions of altered copy numbers. (b) SNPs were grouped in bins of different depth of coverage. SNPs in regions RAF < 0.4 or RAF > 0.6 were excluded from this analysis. When applying the binomial test (purple and blue lines), the frequency of ASB detection increases for higher covered sites, while the same is not true when applying BaalChIP or the iASeq methods. The effect is particularly visible for the FAIRE-seq dataset.

First, for consistency of the analysis, we obtained the set of heterozygous SNPs to test using the BaalChIP QC step. We then compared BaalChIP to the binomial test method with (p=RAF) and without (p=0.5) copy-number correction based on how many of the identified ASB SNPs were in CNA regions. For the comparison to the iASeq method, we computed the estimated posterior probability obtained for SNPs different regions of altered copy-numbers. Figure 5a shows that the binomial test (without RAF correction) and the iASeq methods show a substantial bias towards the increased detection of ASB events in regions altered copy-number, as expected.Both BaalChIP and iASeq employ a beta-binomial distribution to model read count data, which accounts for extra-binomial variability (overdispersion). To check the effect of overdispersion in the ENCODE and FAIRE-seq data we considered only SNPs at regions of normal or close to normal copy numbers (RAF betwen 0.4-0.6) and computed the percentage of detected ASB sites in bins of increasing depth of coverage. The overdispersion is often more accentuated for sites of higher coverage (> 200 pooled total counts), which will create a particularly sensitive scenario for the detection of ASB sites using the binomial distribution. Figure 5b demonstrates clearly the effect of overdispersion in the FAIRE-seq data, particularly visible at sites of high coverage, which shows the advantage of BaalChIP and iASeq over the binomial model. This effect is less pronounced in the ENCODE data, where depth of coverage is overall lower (Figure S16).

These results suggest that for higher read counts random fluctuations in allelic ratios should be modelled with a beta-binomial distribution, rather than a binomial distribution, to reduce the number of false-positive results.

## Discussion

ASB analysis is an important method for identifying putative regulatory SNPs that might have an effect on transcription factor binding and gene expression and might be associated with disease phenotypes. Our method for Bayesian Analysis of Allelic imbalances from ChIP-seq data, called BaalChIP, has been motivated by the need to address the issue of detecting allelic-specific imbalances from ChIP-seq data obtained specifically from cancer cell lines, which frequently carry copy-number alterations.

While allele-specific copy number alterations can be associated with changes in transcript abundance [38] and are possibly implicated in cancer phenotype, they can confound the identification of allele-specific binding at potentially regulatory sites. Traditional methods to identify ASB have not been able to distinguish between effects that are caused by allele specific amplification of binding sites versus the effects caused by allele-specific binding by specific TFs. Our method now allows to distinguish these and will therefore help to better delineate the causal mechanisms of disease.

BaalChIP is a general framework applicable to a wide range of assays. We have applied it to ENCODE ChIP-seq and our own FAIRE-seq samples, and demonstrated the utility of BaalChIP to identify ASB events from cancer genomes. Each of these chromatin assays has its own advantages for the detection of allele-specific binding. ChIP-seq precisely determines the location of specific DNA-associated proteins, while FAIRE-seq identifies broader regions of open chromatin which might be less informative in terms of pinpointing the functional regulatory elements of the genome. On the other hand, the ChIP assay is limited to the availability of high-quality antibodies and only one factor can be tested per experiment. It is still an open question which chromatin assay will be the most informative for understanding allele-specific binding differences.

Cancer cell lines carry frequent copy number alterations. Therefore an important issue is the determination of the background biases in allele frequency observed in regions of altered copy-number. For the ENCODE dataset, we have demonstrated that the use of BAF scores obtained from microarrays could be used to correct for these effects in the allelic ratios, however it is also noteworthy that the microarray technology used by the ENCODE study did not generate BAF scores comprehensively for all possible heterozygous sites, limiting the number of SNPs at which the allele-specificity of each could be assayed. Therefore, for the majority of the samples in the ENCODE case study, it is likely that the regulatory potential of true causal SNPs was not directly assayed. As demonstrated by the analysis of FAIRE-seq data, this issue can be overcome by including genomic DNA-sequencing as control samples to detect allele-specific biases, provided that there is sufficient read coverage at each site.

The identified ASB sites may form the basis for future functional analysis of the genome. Of particular interest, within the FGFR2 gene, rs2981578 has been previously suggested to be the key causal variant. Indeed this SNP displays the highest allelic imbalance in a breast cancer cell line (Table 1).

BaalChIP has been designed for data from cell lines, not tumor samples [39]. Normal contamination and the existence of different clonal subpopulations in tumor samples pose additional challenges and can distort the expected distribution of heterozygous variant allele fractions. In the future, these challenges can potentially be overcome by combining the ideas implemented in BaalChIP with probabilistic methods developed for dissecting genetic heterogeneity and cancer evolution [40].

## Conclusions

In summary, BaalChIP is a rigorous probabilistic method to detect allelic imbalance by correcting for the effect of background allele frequency on the observed ChIP-seq read counts. BaalChIP implements stringent filtering and preprocessing steps and allows the joint analysis of multiple ChIP-seq samples across a single variant. In simulations, BaalChIP outperformed competing approaches, and in an application to 548 ENCODE ChIP-seq and 6 targeted FAIRE-seq samples BaalChIP effectively corrected allele-specific analysis for copy number changes and increased the power to detect putative cis-acting regulatory variants in cancer genomes.

## Methods

### BaalChIP Quality Control and filtering

The QC and filtering steps implemented in the BaalChIP R package are:

1. Keep only reads with MAPQ > 15 and with base quality > 10
2. Within each cell line only consider heterozygous SNPs overlapping transcription factor binding sites identified by ChIP-seq peak calling;
3. Exclude sites susceptible to allelic mapping bias in regions of known problematic read alignment [22, 23, 24];
4. Apply simulation-based filtering to exclude SNPs with intrinsic bias to one of the alleles [6, 17];
5. Consider only SNPs that are represented in all replicated samples after applying all previous filters;
6. Exclude possibly homozygous SNPs where only one allele was observed after pooling ChIP-seq reads from all examined samples [4, 26].

#### Generating allele counts per SNP

For each SNP, the number of reads carrying the reference and alternative alleles are computed using the pileup function and the PileupParam constructor of the Rsamtools package [41]. For each BAM file, BaalChIP only considers heterozygous SNPs overlapping the genomic regions in the corresponding BED files (peaks). Two arguments of the PileupParam constructor can be manipulated by the user: the min_mapq which refers to the minimum “MAPQ” value for an alignment to be included in pileup (reads with mapping quality score lower than this threshold are ignored; default is 15); and the min_base_quality which refers to the minimum quality value for each nucleotide in an alignment (bases at a particular location with base quality lower than this threshold are ignored; default is 10).

#### Data sources for regions of problematic alignments

BaalChIP considers three sets of regions known to be of problematic read alignment, by default: (1) blacklisted regions downloaded from the UCSC Genome Browser (mappability track; hg19, wgEncodeDacMapabilityConsensusExcludable and wgEncodeDukeMapabilityRegionsExcludable tables), (2) non-unique regions selected from DUKE uniqueness mappability track of the UCSC genome browser (hg19, wgEncodeCrgMapabilityAlign50mer table), and (3) collapsed repeat regions downloaded from [24] at the 0.1 % threshold. Sets of filtering regions used in this filtering step are fully customized and additional sets can be added by the user as GenomicRanges objects [42].

#### Simulations to identify SNPs with intrinsic biases

For each heterozygous site, BaalChIP simulates every possible read overlapping the site in four possible combinations — reads carrying the reference allele (plus and minus strand), and reads carrying the alternative allele (plus and minus strand). The simulated reads are constructed based on published methodology [6] (scripts shared by Degner *et al.* upon request) without taking into consideration different qualities at each base in the read or different read depth of coverage. As described by Degner *et al.* these parameters were sufficient to predict the SNPs that show an inherent bias [6].

Reads are then aligned to the reference genome and BAM files are generated using Bowtie version 1.1.1 [43] and Picard tool version 1.47. The pipeline used to generate and align simulated reads can be fully customized with other aligners (e.g. BWA) and it is available under the file name “run_simulations.sh” found in the folder “extra” of the BaalChIP R package. Simulated allelic read counts are computed using the pileup function and the PileupParam constructor of the Rsamtools package [41]. For each SNP, the correct number of reads that should map to the reference and non-reference alleles is known (corresponds to twice the read length for each allele); sites with incorrect number of read alignments are discarded from the analysis.

#### Estimating reference mapping bias

The reference mapping bias is calculated as described in [4, 26]. First, reads are combined across all heterozygous sites that pass the previous QC steps. The expected allelic ratio (REF/TOTAL) for each cell line is then calculated separately for each allele combination. A minimum of 200 sites is required for each category; if there were less, a global estimate was used for that category. For each SNP this computation results in an allelic ratio *μ ∈* (0,1). These allelic ratios are used as part of a prior distribution in the Bayesian model described in the next section.

### BaalChIP Bayesian model description

BaalChIP is designed to infer allelic imbalance from read counts data while integrating copy number and technical bias information. We assume that we are given a set of *N* datasets covering a SNP position. We jointly infer allelic imbalance from signals in all datasets covering the same SNP.

For the *n*th dataset let *d_n_* ∈ ℕ denote the total number of reads covering a SNP position and *a_n_* ∈ ℕ the number of reads reporting the reference allele. BaalChIP models *a_n_* with a beta-binomial distribution, i.e. the binomial distribution in which the probability of success is integrated out given that it follows the beta distribution:

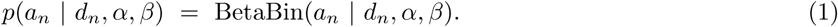

The beta-binomial distribution controls for overdispersion, i.e. the fact that the increased variance in NGS data cannot be captured in the standard binomial model [18]. To gain a more intuitive interpretation of the parameters, we reparametrize *α* and *β* in terms of the precision of the beta-binomial distribution, *Λ*, and the mean probability that a reference read is observed, *θ*:

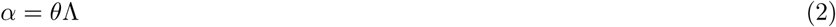

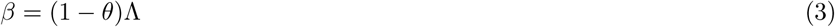

#### Including reference mapping bias

In the next level of our model, we place a beta distribution over *θ*, which allows us to naturally include the technical reference mapping bias:

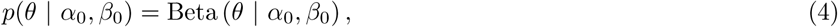

where, as before, we re-parametrise the shape parameters *α*_0_ and *β*_0_ by the mean and variance of the distribution. Denoting the variance by *λ* and using as the mean the reference mapping bias *μ* ∈ (0,1) as discussed above, we get:

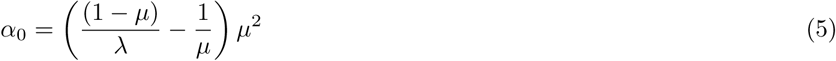

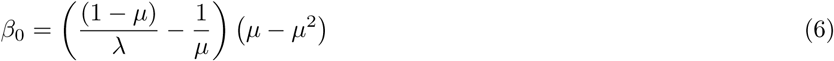

As a consequence of this re-parametrisation, the reference mapping bias has a very intuitive interpretation as the *a priori* mean probability that a read reporting the reference allele is observed.

#### Including reference allele frequency

Next, we reparameterize *θ* as a function of allelic balance ratio *η* and reference allele frequency *ρ*:

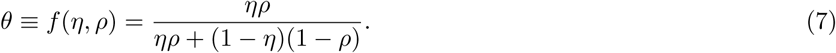

Intuitively, this parameterization implements the Bayes’ rule with the following definitions:

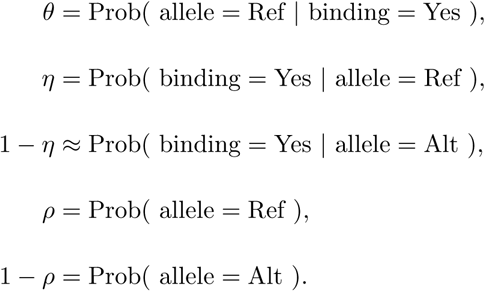

The auxiliary condition on binding status in the definition of *θ* reflects the fact that we only analysed peaks and thus assume that observed ChIP-seq reads are obtained after TF binding. Eq. 7 then formulates a ChIP-seq experiment as a process to obtain the posterior probability of a TF binding to the reference allele. In this model, the allelic balance ratio acts as the likelihood for TF binding to the reference allele and the RAF as the prior.

To better understand how this model works, we will discuss two illustrative scenarios. Example 1: Assuming no copy number alterations, i.e. RAF *ρ* = 0.5, and no allelic imbalance, i.e. *η* = 0.5, then 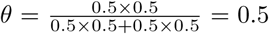, which shows that BaalChIP contains the assumption of previous approaches as a special case. Example 2: In the event of LOH on the alternative allele, RAF *ρ* =1, then irrespective of the allelic balance ratio, all observed reads will report the referenceallele, which is reflected by 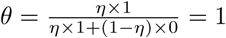

#### Posterior distribution of allelic imbalance

To rigorously represent the uncertainty in the data, we adopt a full Bayesian approach targeting the posterior distribution of allelic balance ratio *η*. We use the fact that the change of variables in Eq. 7 and the prior on *θ* directly define a prior over *η*, namely

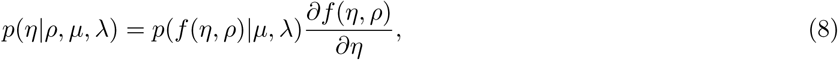

and write the posterior of *η* as follows:

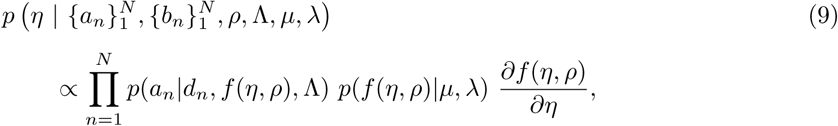

where

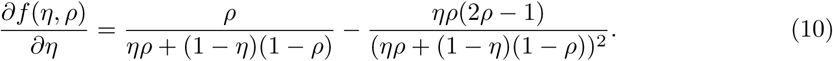

#### Inference of allelic imbalance

We use the Metropolis-Hastings algorithm with random walk proposal to approach this distribution. In practice, we fix the Λ = 1000 and λ = 0.05, which in our experience results in robust performance across a wide range of simulated and real datasets. Allelic imbalance calling is based on the highest posterior density (HPD) interval, which is constructed from the MCMC trace of *η* as the shortest interval containing 95% of sampled values. Allelic imbalance is called, if the HPD does not contain the value 0.5, which is a rigorous way to decide that the data cannot be explained well by balanced alleles.

### *In silico* validations

To thoroughly test the performance of allelic imbalance calling performance of BaalChIP, read counts data are generated using a wide range of parameter settings. Specifically, we varied the number *N* of data sets from 1 to 45, *ρ* from 0.1 to 0.9 in steps of size 0.1, *d* from 1 to 100 in steps of size 7, and 1000 values of *η* are evenly sampled from 0.1 to 0.9. These settings result in a set of 6,075,000 SNPs. The detailed simulation protocol is as follows:

For each *n* ∈ [1,…, N] and *ρ* ∈ [0.1,…, 0.9] and *η* ∈ [0.1,…, 0.9]:

1. Draw the number of available reference alleles:

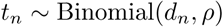
2. Draw the read counts reporting reference allele:

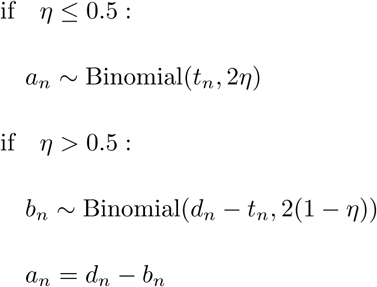

The performance is measured by ROC curves. The true allelic imbalance is determine if the true *η* is outside the interval [0.45, 0.55].

### Baseline methods

We compared BaalChIP against two baseline approaches: binomial test and iASeq [10]. Two-sided binomial tests are performed with the R function binom.test. To correct for reference mapping bias, the null hypothesis on the probability of success is set to be the previously estimated reference mapping bias (RAF), instead of 0.5. We pooled data from all ChIP-seq samples for each analyzed cell line to maximize power. For the analysis of real data, binomial test p-values were corrected for multiple testing using the false discovery rate (FDR) threshold of 0.01. FDR was calculated with the p.adjust function in R. For the simulated data, the ROC curves for binomial test are constructed based on the distance between reference mapping bias and the mean of 95% confidence intervals for the probability of success. All iASseq analyses are performed with the function iASeqmotif where the number of non-null motif is set to be a vector [1, 2,…, 5], the maximun number of iterations is 300 and the tolerance level of error is 0.001. The ROC curves for iASseq are constructed with bestmotif$p.post which is the posterior probability for each SNP being allele-specific event.

### Cell culture

MDA-MB-134 and T-47D human breast cancer cells were cultured in RPMI (Invitrogen) supplemented with 10% FBS and antibiotics. All cells were maintained at 37C, 5% CO2. All cell lines were from the CRUK Cambridge Institute biorepository collection. Cell lines were authenticated by short tandem repeat genotyping using the GenePrint 10 (Promega) system and confirmed to be mycoplasma free.

### FAIRE and gDNA purification

FAIRE stands for ‘Formaldehyde assisted isolation of regulatory elements’ and was performed as previously described [35] with minor adaptations. Briefly, cells were fixed for 10 mins in 1% formaldehyde in FCS-free medium, washed and frozen. Nuclei from MDA-MB-134 (3 × 10^7^ cells/tube) and T-47D (1.5 × 10^7^ cells/tube) were isolated and sonicated using 300*μ*l volumes in 1.5ml eppendorfs, using a Diagenode Bioruptor. Sonication was performed for 20 cycles of 30s on/off at the “high” setting. The sonicate was subjected to three consecutive phenol-chloroform-isoamyl alcohol (25:24:1) extractions and reverse cross-linked overnight. DNA was purified by ethanol precipitation and quantified by Quanti-iT. Genomic DNA was isolated using Qiagen DNeasy Blood and tissue Kit according to protocol.

### Agilent SureSelect amplification, library preparation and sequencing

DNA fragments were prepared for sequencing using the recommended protocol available for SureSelect-Illumina sequencing. 69 known breast cancer risk tagging SNPs were retrieved from [36]. SNAP Proxy Search was used to find SNPs correlated with the 69 tagging SNPs (r2 > 0.6 for 59 of the risk loci and r2 > 0.8 for 8 of the risk loci, within a distance of 500bp) using 1000 Genomes pilot 1 data. Genomic intervals were defined by the left-most to the right-most SNP in each LD block with an additional 400bp of flanking regions. An additional set of 7 SNPs with 500bp of flanking sequences were added manually to include the CCND1 (rs75915166, rs494406), MAP3K1 (rs10461612, rs112497245, rs17432750) and TERT gene regions (rs2736107, rs7705526). In total, we targeted 69 non-overlapping regions comprising 4.93Mb using SureSelect method (Table S8). DNA obtained from SureSelect solution-based sequence capture was subjected to Illumina HiSeq paired-end sequencing (Illumina). Paired-end Sequencing was performed according to manufacturer’s protocols.

### Pre-processing ENCODE samples

We used publicly available ENCODE ChIP-seq and genotype datasets for a total of 548 samples representing 271 different experiments. We included 8 cancer and 6 non-cancer cell lines representing different tissues. Table S3 shows a summary of all tissues, cell lines and number of experiments included in this study. The ChIP-seq data was downloaded as reads mapped to the hg19 genome (BAM files) and correspondent peak calling files (BED files). Accession numbers of all public ChIP-seq datasets used in this study are provided in Table S2. Duplicated reads were marked using Picard tool version 1.47. ChIP-seq peak files were merged between replicates using HOMER version 5.4 mergePeaks with the default option–d given to only consider peak ranges that overlapped for all replicates. Heterozygous SNPs and BAF tracks were retrieved from the UCSC Genome browser (hg19, wgEncodeHaibGenotype track, wgEncodeHaibGenotypeBalleleSnp2015-03-04.tsv and wgEncodeHaibGenotypeGtypeSnp2014-09-15.tsv files). The initial number of genotyped SNPs in the ENCODE files is 1.2 million SNPs. We only considered SNPs listed in dbSNP (version 137 [44]). Homozygous SNPs (i.e., BAF > 0.9 or < 0.1) and SNPs with missing BAF scores were removed from the BAF tracks. BAF scores were converted to Reference Allele Frequency (RAF) scores using the information of A and B alleles in the Manifest file for the Illumina Human1M-Duo BeadChip (v3.0) downloaded from http://support.illumina.com/array/array_kits/human1m-duo_dna_analysis_kit/downloads.html.

### Pre-processing FAIRE-seq samples

For sequence data from all FAIRE-seq and control samples, sequences were aligned to the human reference genome (GRCh37) using BWA version 0.7.12 [45]. Duplicates were removed using Picard Tool version 1.47 and overlapping reads were clipped using clipOverlap tool from bamUtil repository version 1.0.14 with default parameters. SNPs were identified from gDNA sampes using the Genome Analysis ToolKit 3.4-46 software [46] across all gDNA samples simultaneously. As per GATK Best Practices recommendations [47, 48], duplicated reads were removed and local realignment and base quality recalibration were employed prior to SNP calling. Called SNPs were filtered using a hard filtering criteria (QD < 2.0, FS > 60.0, MQ < 30.0, MQRankSum < -12.5, ReadPosRankSum < -8.0).

### Applying BaalChIP to ENCODE and FAIRE-seq samples

To ensure a reliable set of heterozygous SNPs we applied the BaalChIP (version 0.1.9) quality control step and considered only uniquely mapping reads with MAQ > 15 and base call quality > 10. For the ENCODE dataset we removed from the analysis sets of SNPs based on the default six QC filters implemented within the BaalChIP pipeline. The FAIRE-seq samples contained paired-end sequenced reads of 125 bp. Since longer and paired-end reads reduce uncertainty of read alignment we did not consider two of the QC filters that are more relevant for shorter read lengths (of less than 50bp): the unique mappability and intrinsic bias filters. BaalChIP (version 0.1.9) ASB Bayesian analysis was performed with the default parameters and options [21].

### Consistency of allelic ratios

Consistency of the allelic ratios observed at ASB SNPs was analysed between (i) between replicates: pairs of replicated samples, (ii) within cell lines: pairs of ChIP-seq datasets from different proteins (pooled replicate data) in the same cell line (iii) between cell lines: pairs of cells (pooled data for each cell line). Correlations were calculated with Spearman correlation and required at least 15 shared ASB sites.

### SNP annotations relative to genes

To annotate ASB SNPs with respect to gene annotations — 5’ UTR, 3’ UTR, promoter, spliceSite, coding, intron or intergenic — we used the VariantAnnotaion (version 1.8.13) package in R. To obtain the list of known genes and coordinates, we used the UCSC genome browser known gene annotation obtained from the TxDb.Hsapiens.UCSC.hg19.knownGene (version 2.6.2) library in R [20].

### Overlap with predicted enhancers

To determine if ASB SNPs were enriched in any of the putative enhancer regions, we calculated the overlap of intergenic ASB SNPs in putative enhancer regions. The significance of the observed overlap was determined by a *χ*^2^ test by comparing the fraction of ASB SNPs in putative enhancer regions with the fraction of non-ASB heterozygous SNPs in putative enhancer regions. Putative enhancer regions were retrieved for 6 cell lines: GM12878, H1hESC, HeLa, HepG2, K562, A549 based on predicted weak and strong enhancer sites and/or H3K27ac and H3K4me1 chromatin marks. Enhancer site predictions were retrieved from human segmentations previously generated based on ENCODE data using Segway (downloaded from wgEncodeAwgSegmentation UCSC genome browser track).

### TF motif disruption analysis

In order to annotate the potential regulatory effects of the tested SNPs on transcription factor binding site (TFBS) motifs, the publically available HaploReg v2 database (accessible at http://www.broadinstitute.org/mammals/haploreg/haploreg_v2.php) was used. Haploreg calculates allele-specific changes in the logodds (LOD) scores for position weight matrices (PWMs) of a regulatory motif based on a library of PWMs constructed from TRANSFAC, JASPAR, and and protein-binding microarray (PBM) experiments [49]. The magnitude of the change in binding affinity was calculated as the absolute difference (DELTA) of LOD scores (DELTA = LOD(ref) — LOD(alt)). The Kolmogorov-Smirnov test was used to compare the distributions of the absolute DELTA scores of motif disrupting ASB SNPs and all tested motif disrupting SNPs. To determine which TFBS-motifs ASB SNPs were more likely to disrupt, SNPs were grouped according to DNA-binding proteins as identified by ChIP-seq peaks. For each group, a Fisher exact test was used to identify TFBS-motifs that were more significantly disrupted by ASB SNPs when compared to non-ASB SNPs.

## Availability of data and materials

ENCODE datasets used in this study are publicly available in the curated Gene Expression Omnibus (GEO) database. Accession numbers of all public ChIP-seq datasets used in this study are provided in Table S2. FAIRE-seq datasets used in this study are included in the GEO database (GEO accession number, GSE85261)

## Author’s contributions

IS, WL and KY designed and developed the Baal-ChIP statistical framework. IS and WL implemented BaalChIP package in R. IS analyzed the data. WL preformed simulation studies. KM and MOR designed and preformed FAIRE-seq experiments. BAJP, FM and KBM supervised the analysis. IS, WL, KY, KBM and FM wrote the manuscript. FM and KBM are co-corresponding authors.

## Acknowledgments

We thank Thomas Carroll, Gordon Brown and Jing Su (CRUK Cambridge Institute, University of Cambridge) for suggestions and advice on ChIP-seq data analysis.

## Supplementary Material

**Figure S1.**
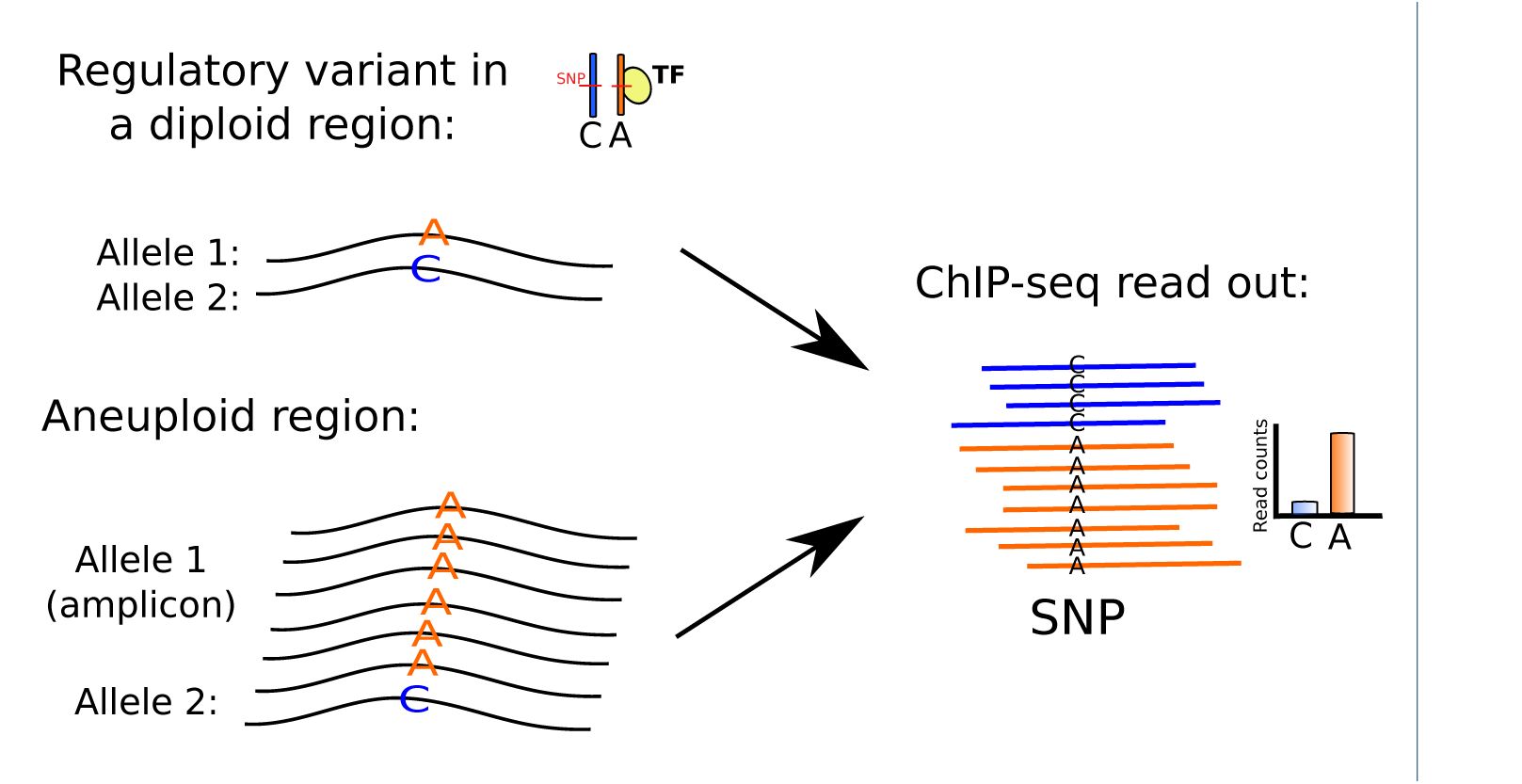
Example illustration of a ChIP-seq read out at a DNA-binding site when a true regulatory difference exists between two alleles (A allele depicted in orange and C allele depicted in blue), and when there is no regulatory difference between the two alleles. In a diploid sample, the protein (yellow circle) binds preferentially to the A allele. Here the regulatory effect is observed as an imbalance in the allelic ratios obtained from ChIP-seq read counts, with a higher number of reads carrying the A allele. In the presence of allele-specific copy number aberrations such as an amplicon affecting the A allele, the direct ChIP-seq readout may reflect the relative presence of the alleles, rather than the regulatory effect of the single nucleotide variant. Consequently, in the presence of copy number changes, ChIP-seq allelic ratios are not sufficient to uncover true cis-regulatory effects.

**Figure S2.**
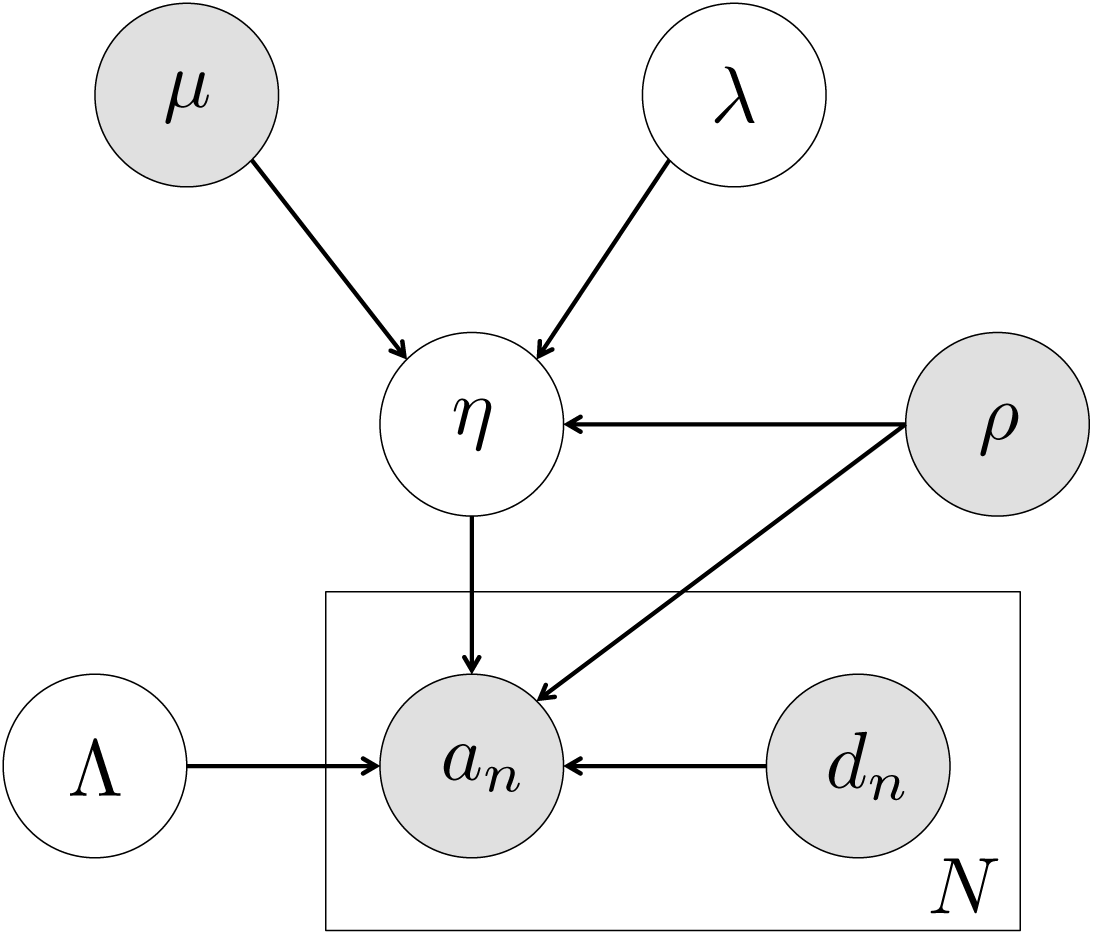
Graphical model of BaalChIP. The gray circles represent given variables and white circles are unknown variables. The direction of arrows indicates statistical dependency in the model. The observed reference allele read count is *a_n_,* total read counts is *d_n_. n* is the index of given factor (e.g. transcription factor) binding to the region of the SNP of interest. The rectangular plane and the capital *N* indicates that there are *N* factors binding to the SNP. The primary variable of interest is *η* representing allelic balance ratio. *Λ* is the precision parameter. The reference allele frequency at the SNP is *ρ. d_n_,η,ρ,Λ* are the parameters of a beta-binomial distribution that governs the uncertainty in *a_n_.* Finally, the reference mapping bias *μ* and λ are the mean and variance of a beta distribution which serves as a prior over η.

**Figure S3.**
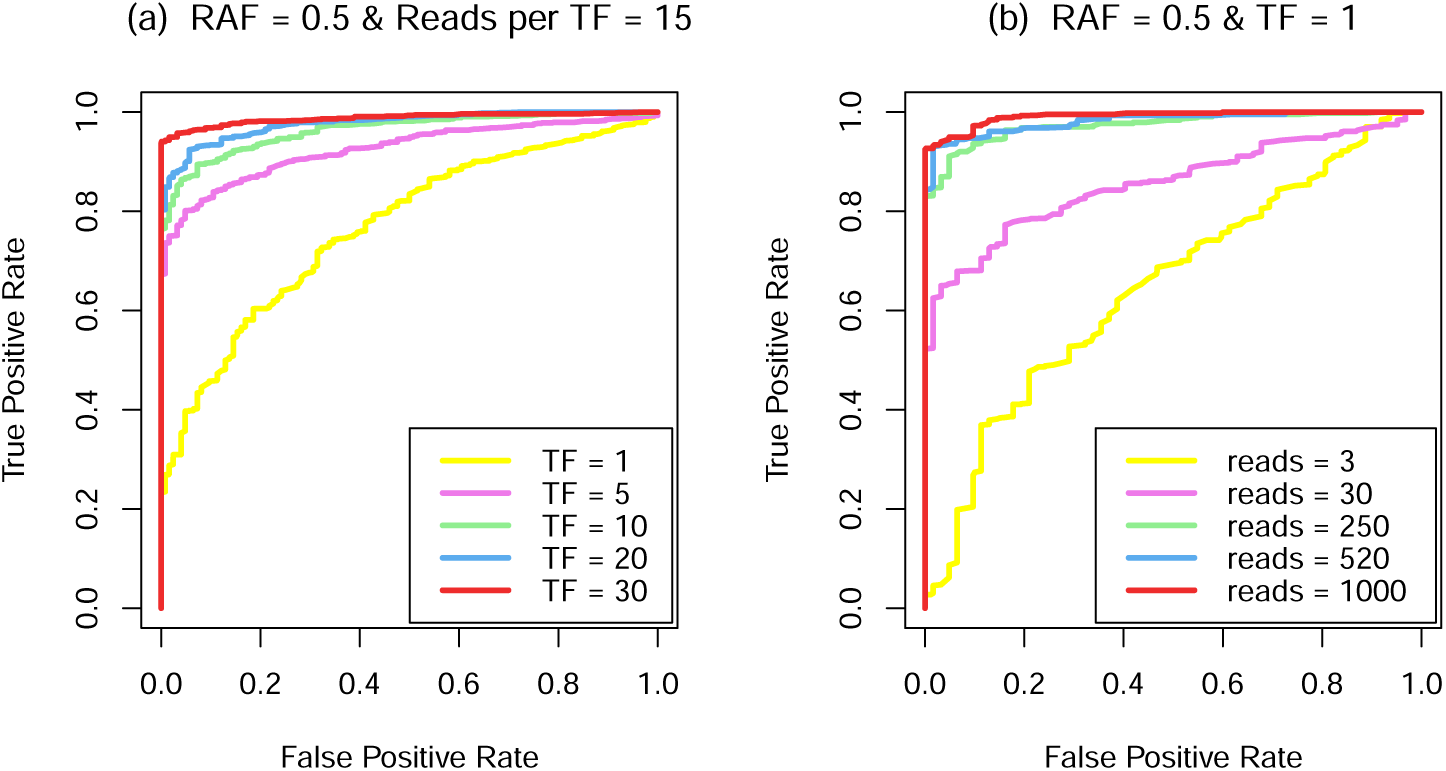
BaalChIP simulation performance with varying number of transcription factors (a) and read depth (b).

**Figure S4.**
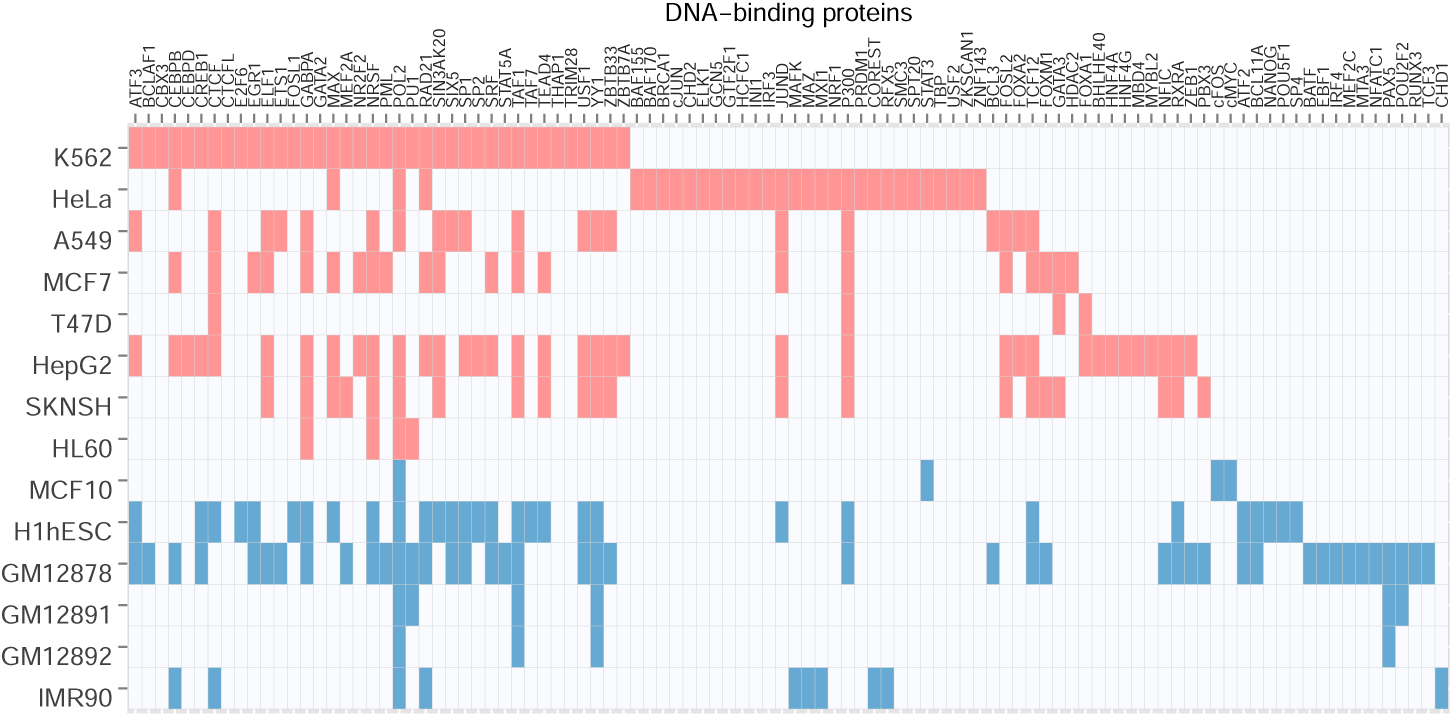
Data matrix summarizing the ENCODE samples used in this study for allele-specific binding analyses. More detailed description including assession numbers can be found in Table S2 and Table S3.

**Figure S5.**
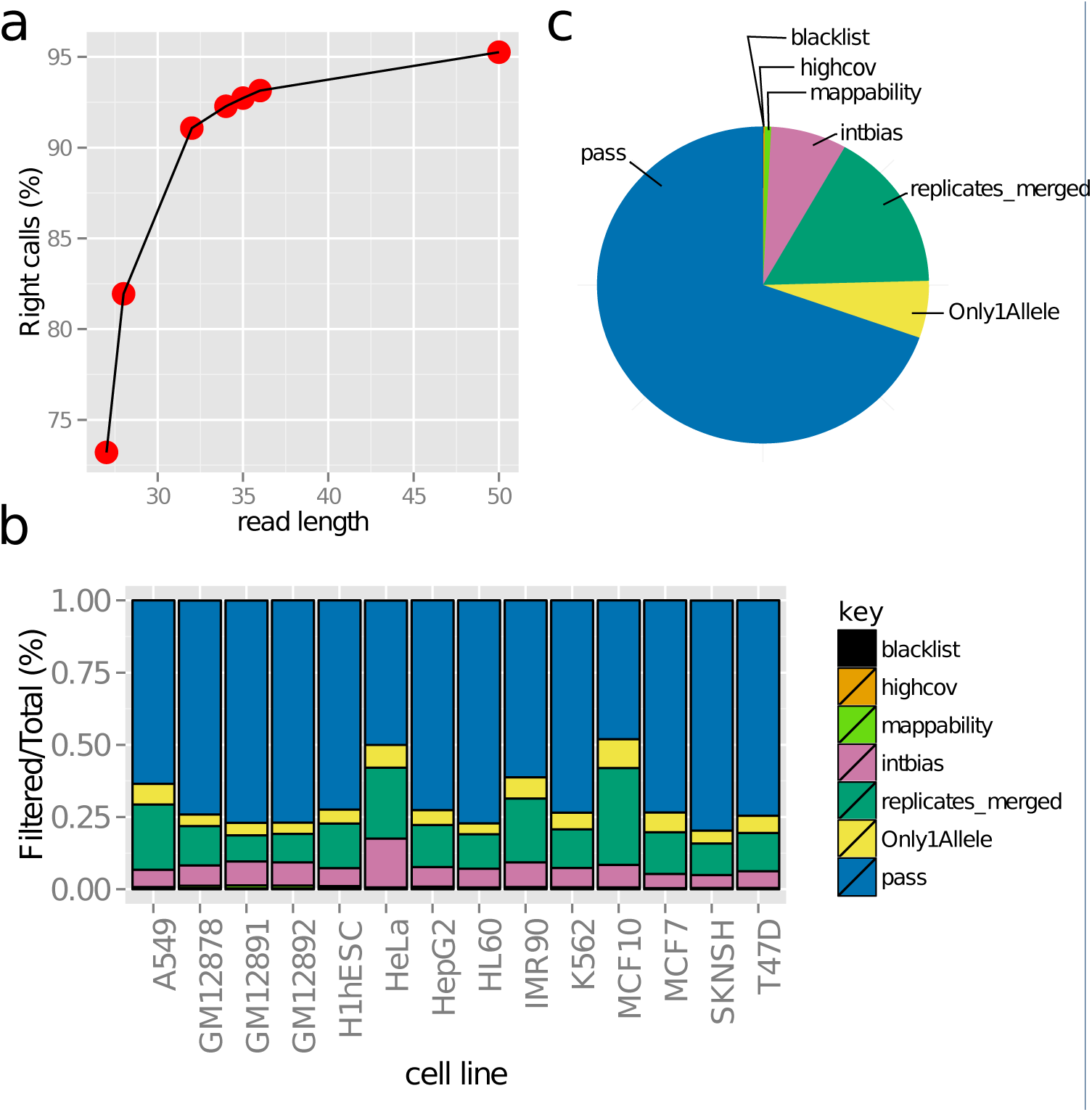
BaalChIP SNP filtering. (a) Percentage of SNPs with the correct number of aligned reads based on simulations of reads of different read lengths (28 to 50 mer). The percentage of correct calls increased with read length. (b) Proportion of SNPs that were filtered out in each filtering step. (b) The average proportion of SNPs that were filtered out in each filtering step. Average was calculated accross all 14 ENCODE cell lines.

**Figure S6.**
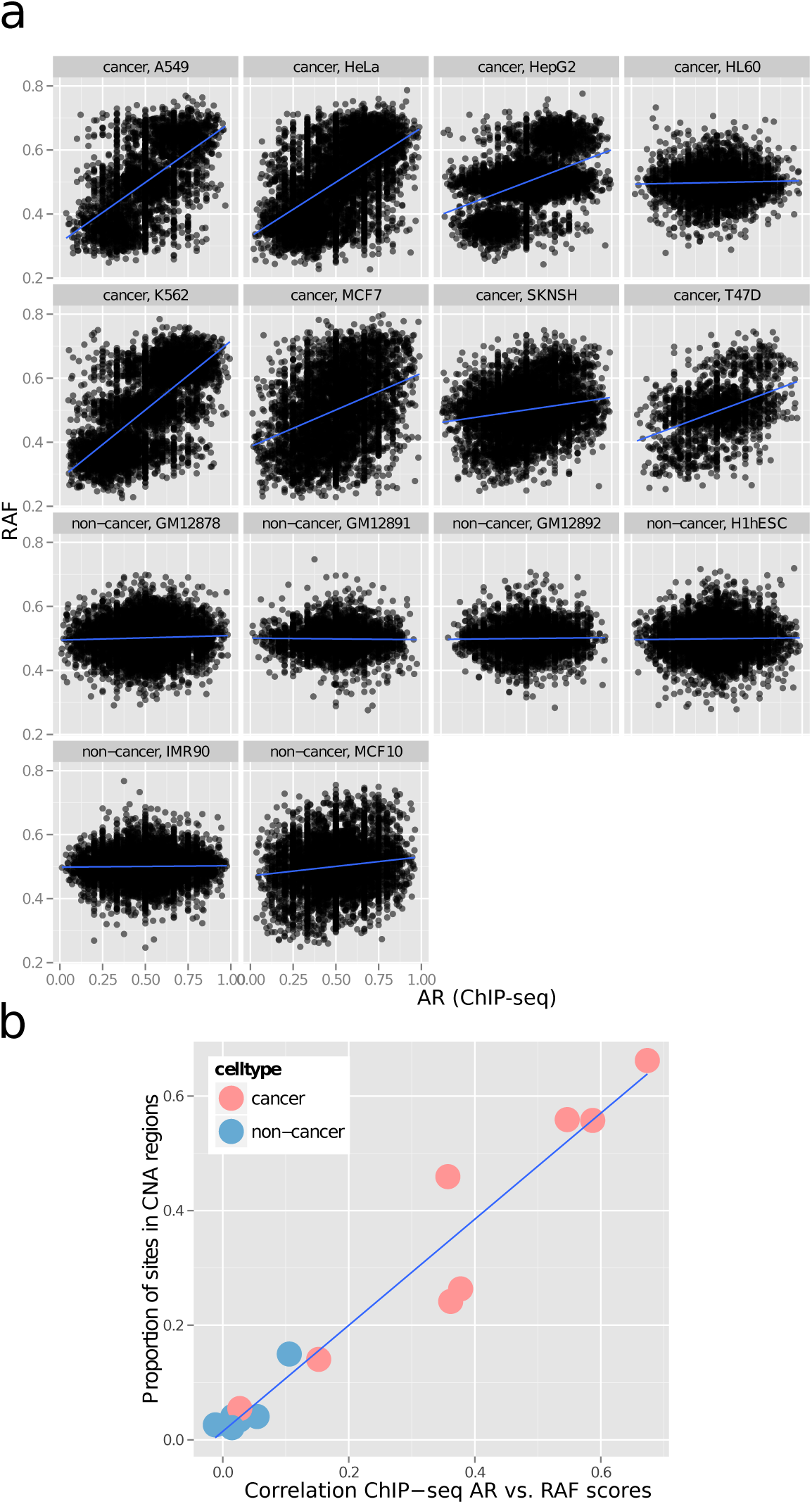
Correlation between the BAF values and the ChIP-seq allelic ratios (AR). (a) Scatter plots showing the correlation between RAF and AR at each SNP, for all considered cell lines of the ENCODE dataset. RAF corresponds to the BAF value with respect to the reference allele (RAF is equal to BAF if the reference allele corresponds to the B allele; RAF is equal to 1-BAF if the reference allele corresponds to the A allele). The blue line shows the fitted linear regression line. (b) Positive relationship between the Spearman correlation coefficient (obtained from the correlation of AR with RAF scores) and the proportion of sites in copy-number altered (CNA) regions. Each dot in the scatter plot corresponds to a cell line. Sites in copy number changed regions correspond to heterozygous SNPs with BAF score < 0.4 or BAF > 0.6.

**Figure S7.**
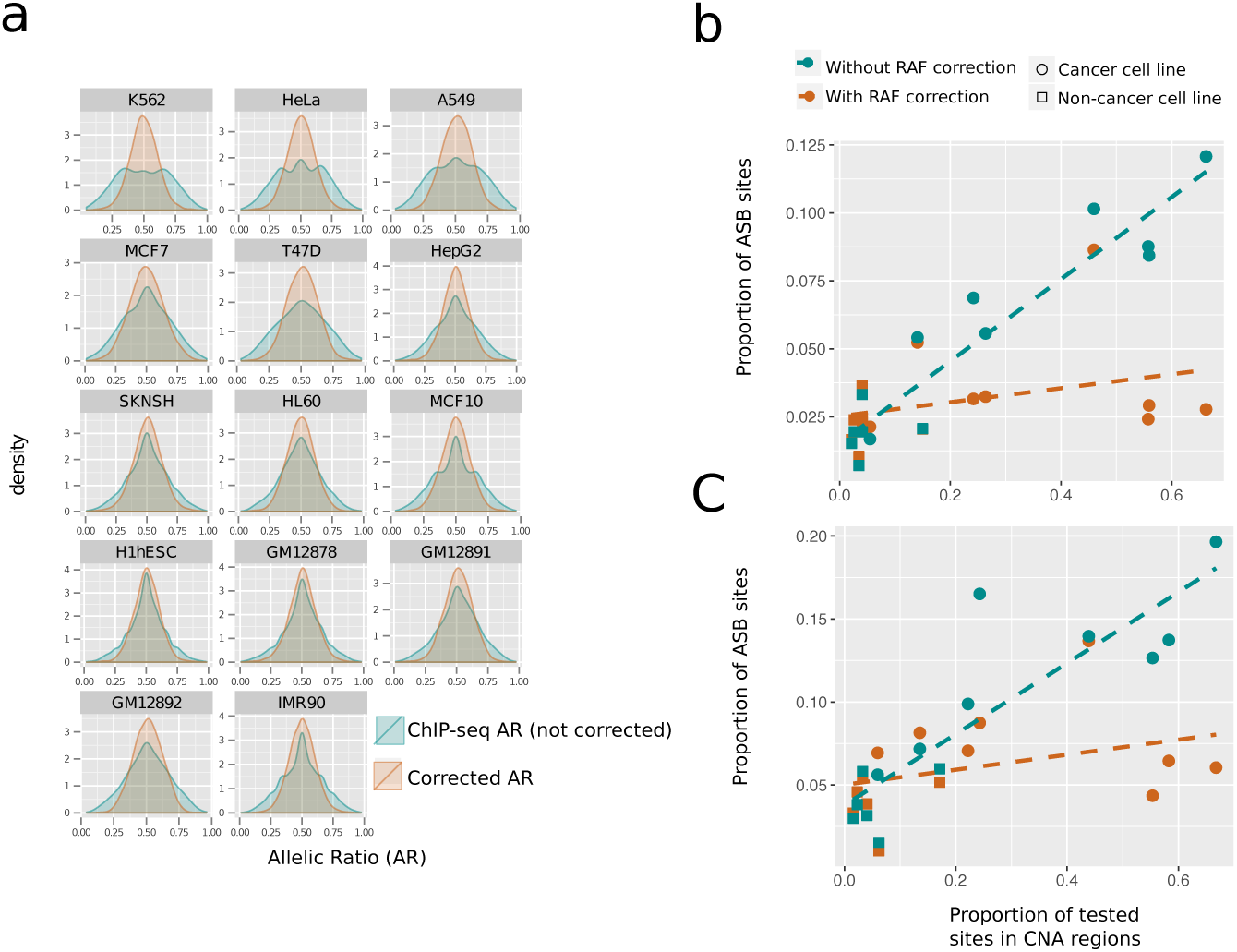
Effects of BaalChIP adjustment of the allelic ratios (AR) after correcting for the background Reference Allele Frequency (RAF) bias. (a) Density plots showing the distribution of allelic ratios before (green) and after (orange) BaalChIP correction. The ARs before correction were calculated directly from the ChIP-seq data and refer to the number of reads in the reference allele divided by the total number of reads. The corrected AR values were estimated by BaalChIP model after taking into account the RAF bias. The adjustment of AR is more dramatic in cancer cell lines. (b) Proportion of ASB sites detected with (orange) and without (green) RAF correction as a function of the proportion of sites within copy-number altered (CNA) regions. Each dot refers to a cell line. Sites in regions of copy number change refer to any SNP with a BAF score higher than 0.6 or lower than 0.4. Cancer cell lines have a higher proportion of SNPs in CNA regions and benefit more from BaalChIP correction. (c) Same as (b) but after selecting SNPs with 30X-40X coverage.

**Figure S8.**
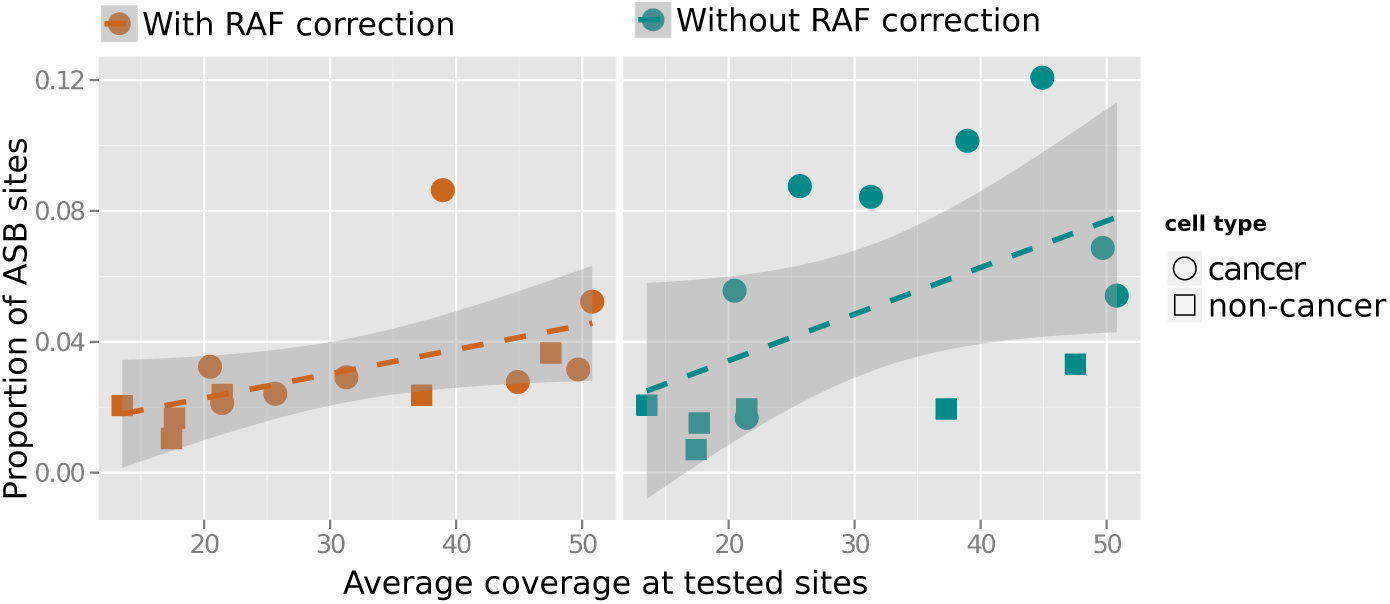
Proportion of identified ASB SNPs per cell line. Correlation between the proportion of identified ASB SNPs and the average coverage at all assayed heterozygous SNPs. The general correlation is expected due to the higher power to detect ASB when coverage is high.

**Figure S9.**
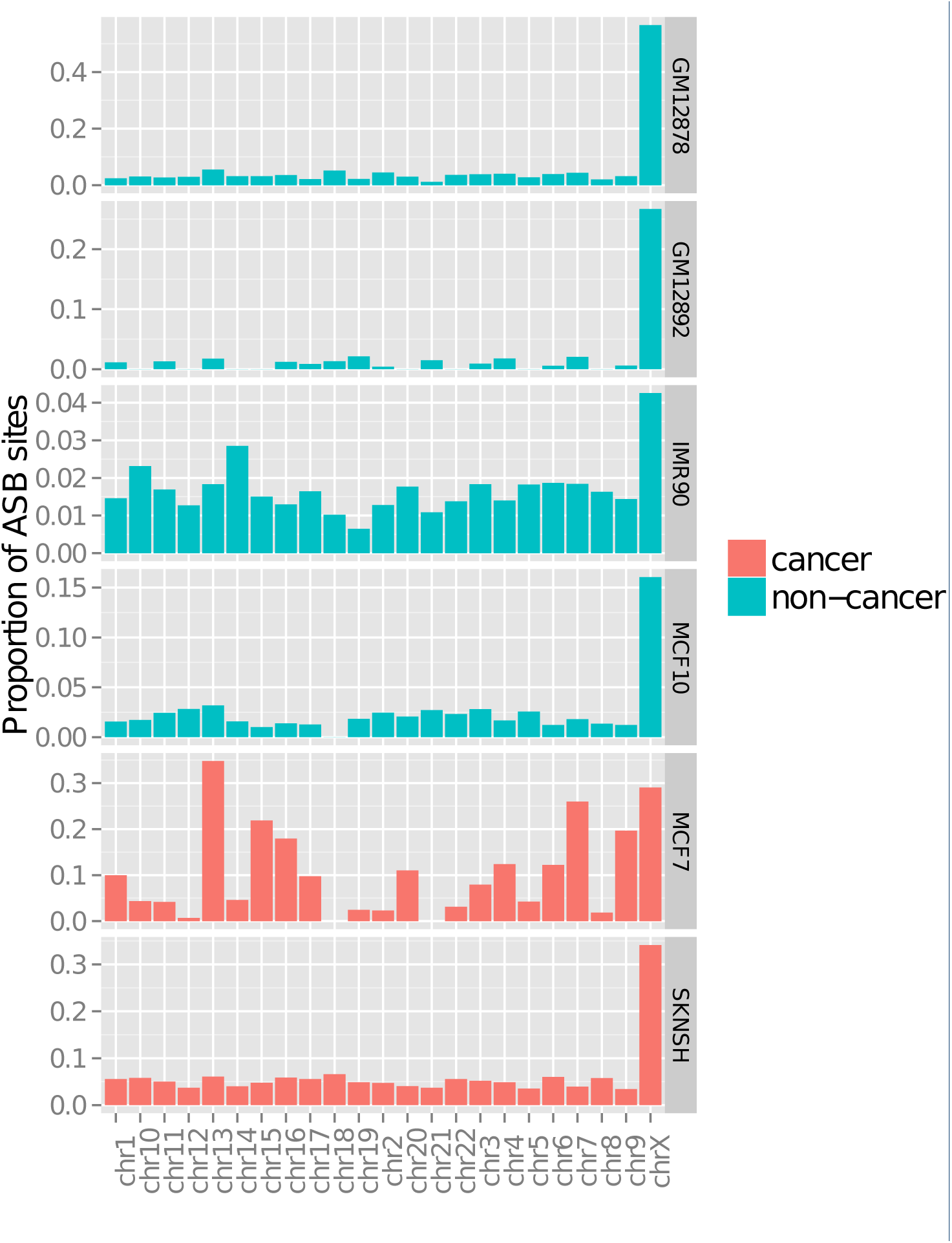
Higher rates of ASB on chromosome X of female cell lines than in autosomal chromosomes.

**Figure S10.**
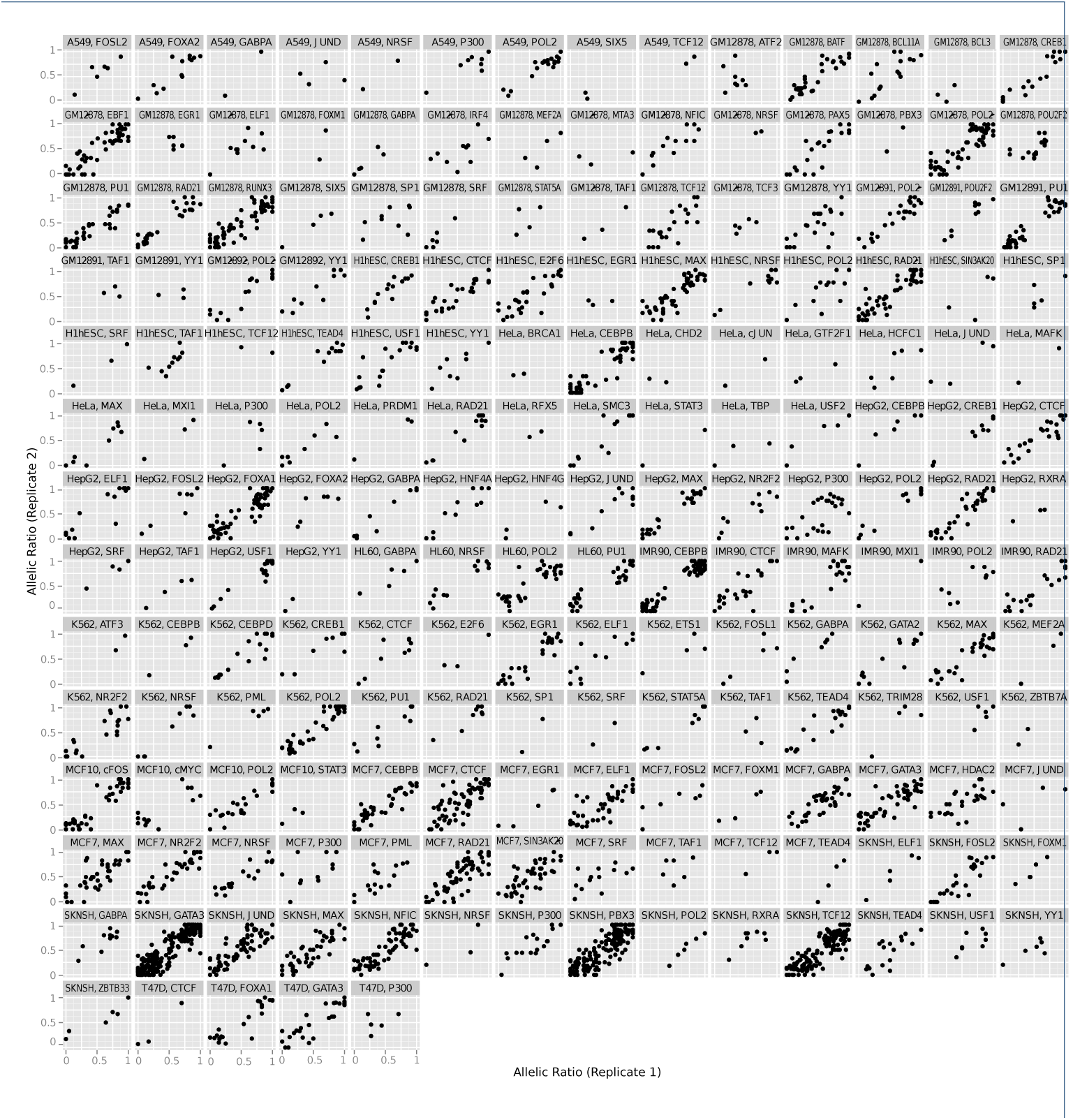
Consistency of ASB across biological replicates. Each dot in the scatter plot represents the allelic ratio at a single SNP obtained for two replicated samples from the same ChIP-seq experiment.

**Figure S11.**
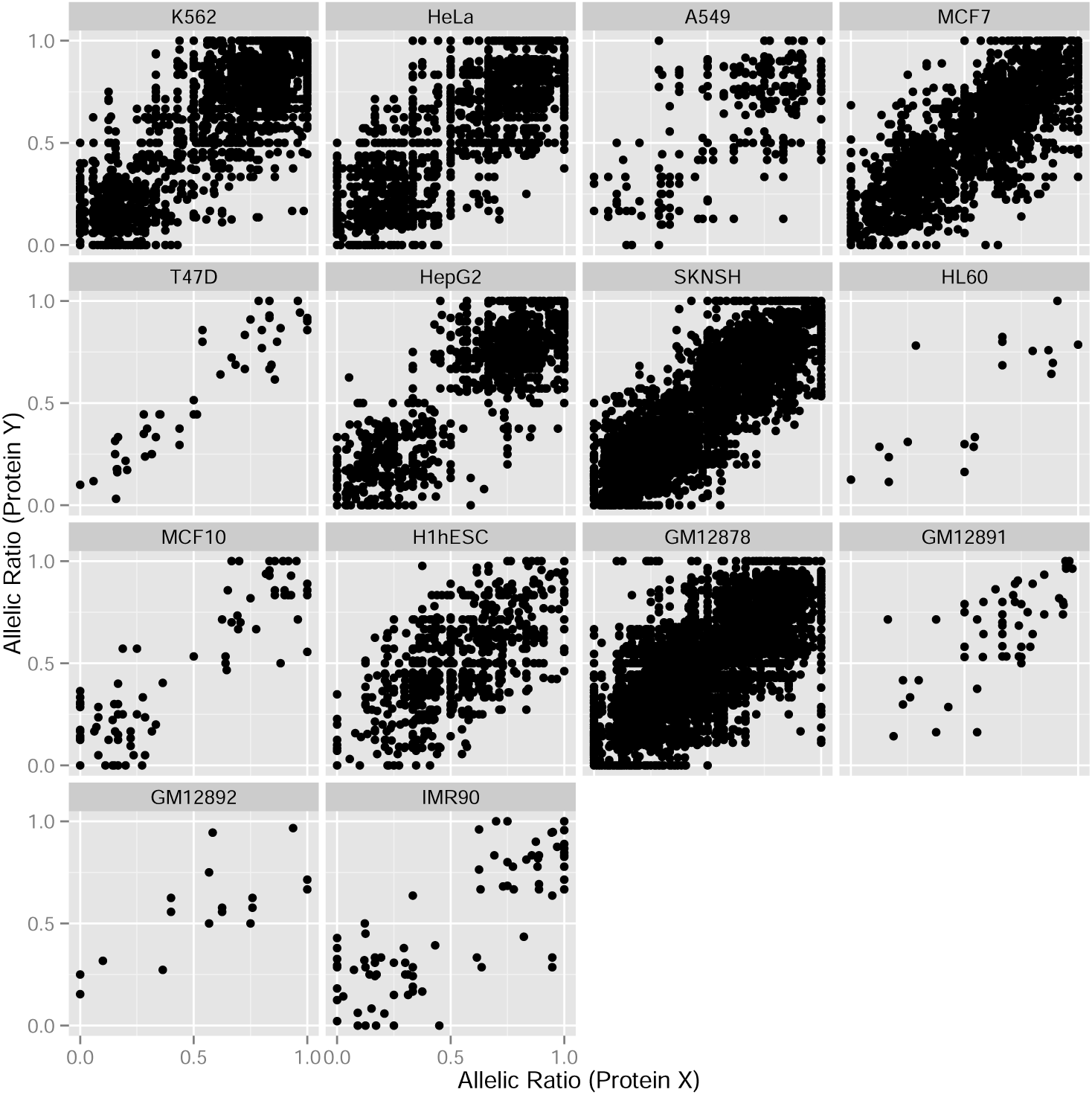
Consistency of ASB across pairs of different assayed proteins (Protein “X” and “Y”), within the same cell line. Each dot in the scatter plot represents the allelic ratio at a single SNP obtained for distinct ChIP-seq experiments in the same cell line (i.e distinct ChIPed proteins bound at the same site).

**Figure S12.**
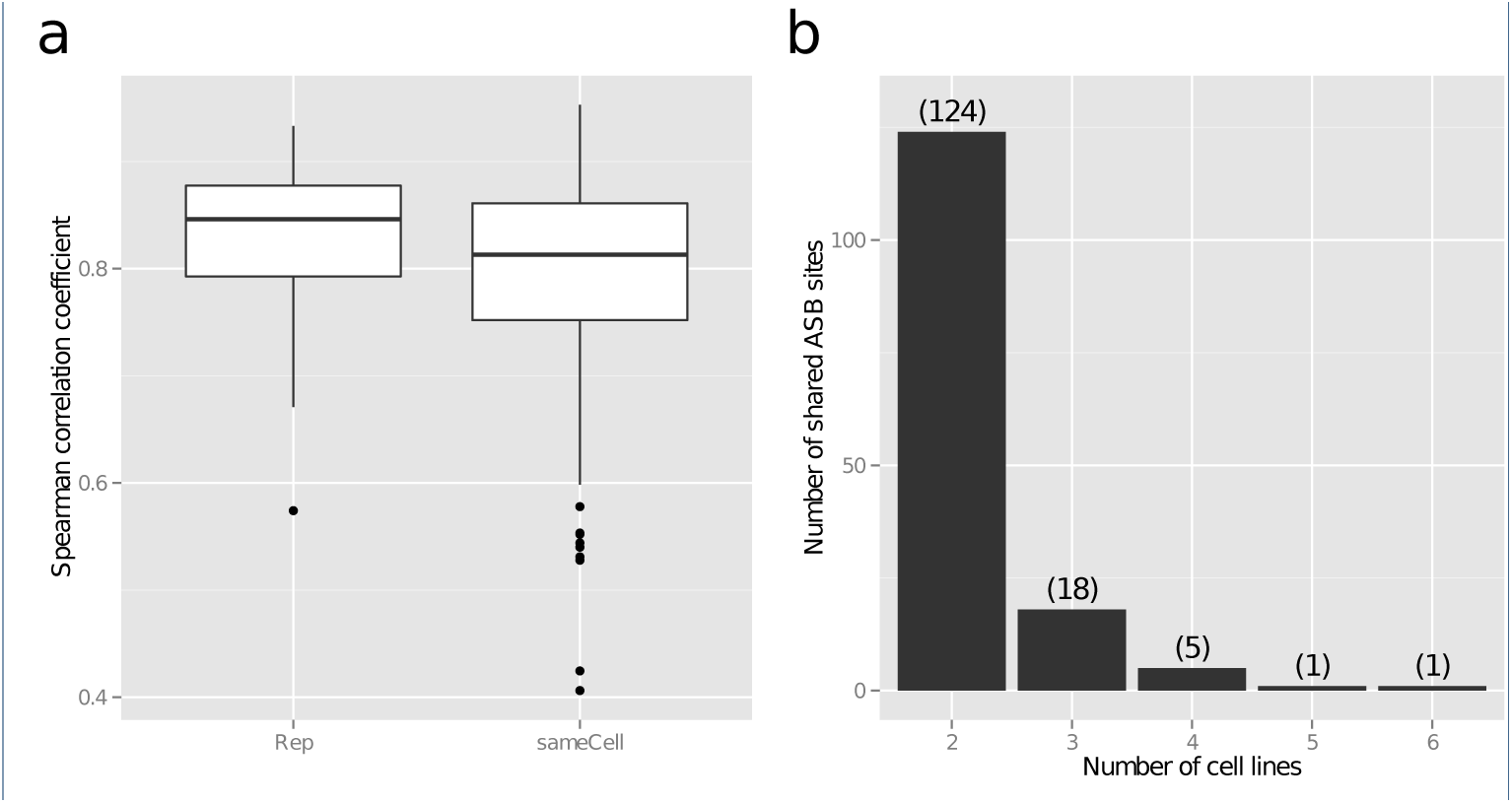
(a) Distribution of Spearman correlation coefficients obtained for the correlation of allelic ratios obtained for pairs of biological replicates (Rep), and between distinct proteins bound at the same site (sameCell). The allelic ratios are highly correlated in both scenarios. (b) Number of shared ASB SNPs across cell lines. Numbers in parenthesis show the number of ASB SNPs shared between 2 to 6 cell lines. 149 ASB SNPs (124+18+5+1+1) were concomitantly detected in two or more cell lines.

**Figure S13.**
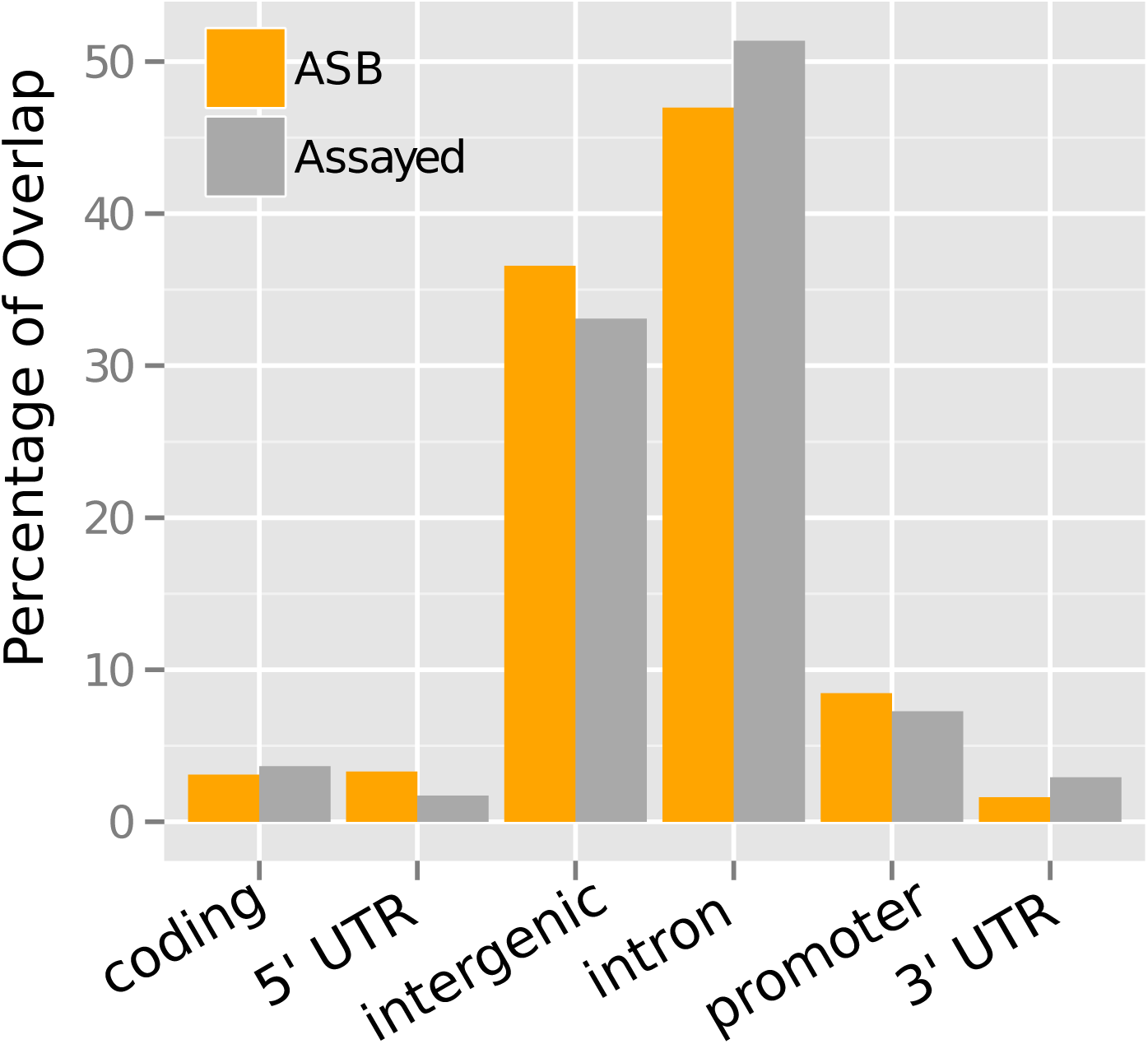
Percentage of ASB SNPs overlapping gene annotations (5’ UTR, 3’ UTR, coding, intergenic, intron or promoter). The largest proportion of ASB SNPs was found in introns and intergenic regions. A similar distribution pattern is observed when considering all assayed heterozygous SNPs.

**Figure S14.**
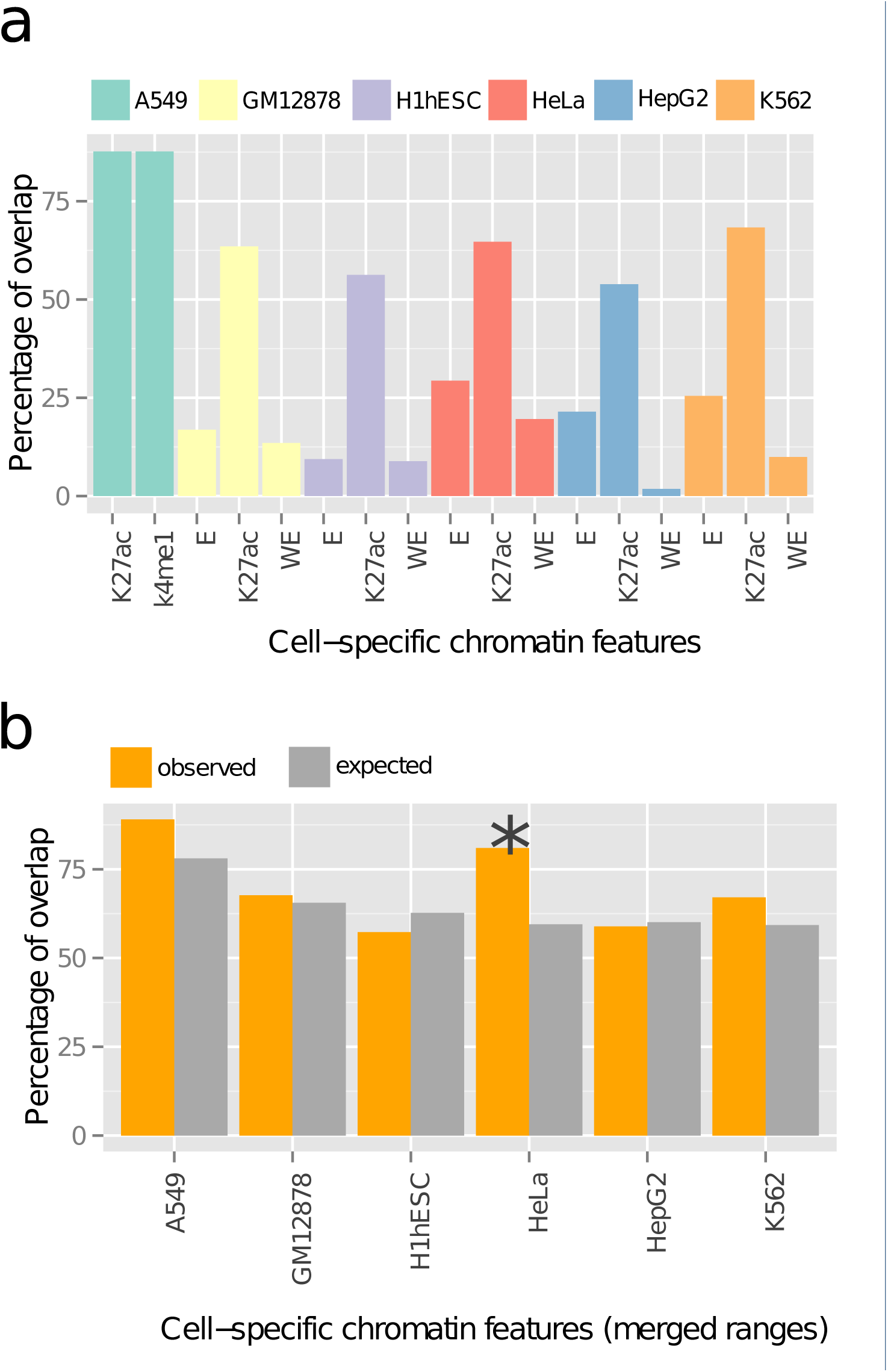
Percentage of ASB SNPs overlapping putative enhancer regions from the matched cell line. (a) Putative enhancer regions were retrieved from publicly available ENCODE datasets on H3K27ac and H3K4me1 modified regions, and previously predicted weak enhancers (WE) and strong enhancer (E) sites obtained from Segway chromatin state annotations of the ENCODE data. (b) Comparison of the observed and expected number of ASB SNPs that map to enhancer regions. A significant difference (*) was only detected in HeLa cells.

**Figure S15.**
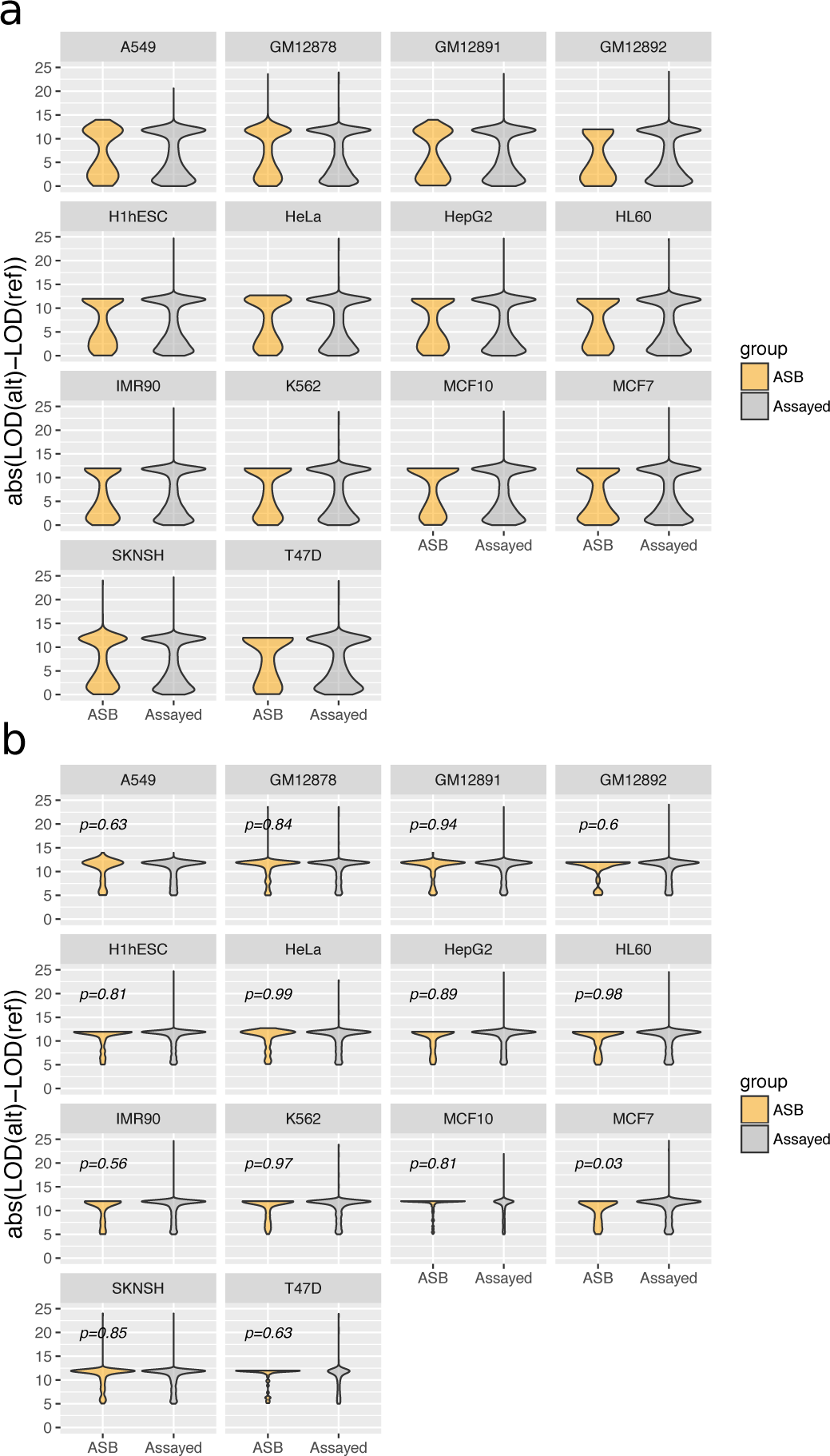
Distribution of absolute DELTA scores (deltaLOD) of motif-disrupting SNPs with (a) deltaLOD > 0 and (b) deltaLOD > 5. The DELTA score corresponds to the change in the PWM score between SNP alleles (DELTA = LOD(ref) — LOD(alt)). P-values correspond to the Kolmogorov-Smirnov test when comparing the distribution of DELTA scores between motif-disrupting ASB SNPs with all motif-disrupting SNPs.

**Figure S16.**
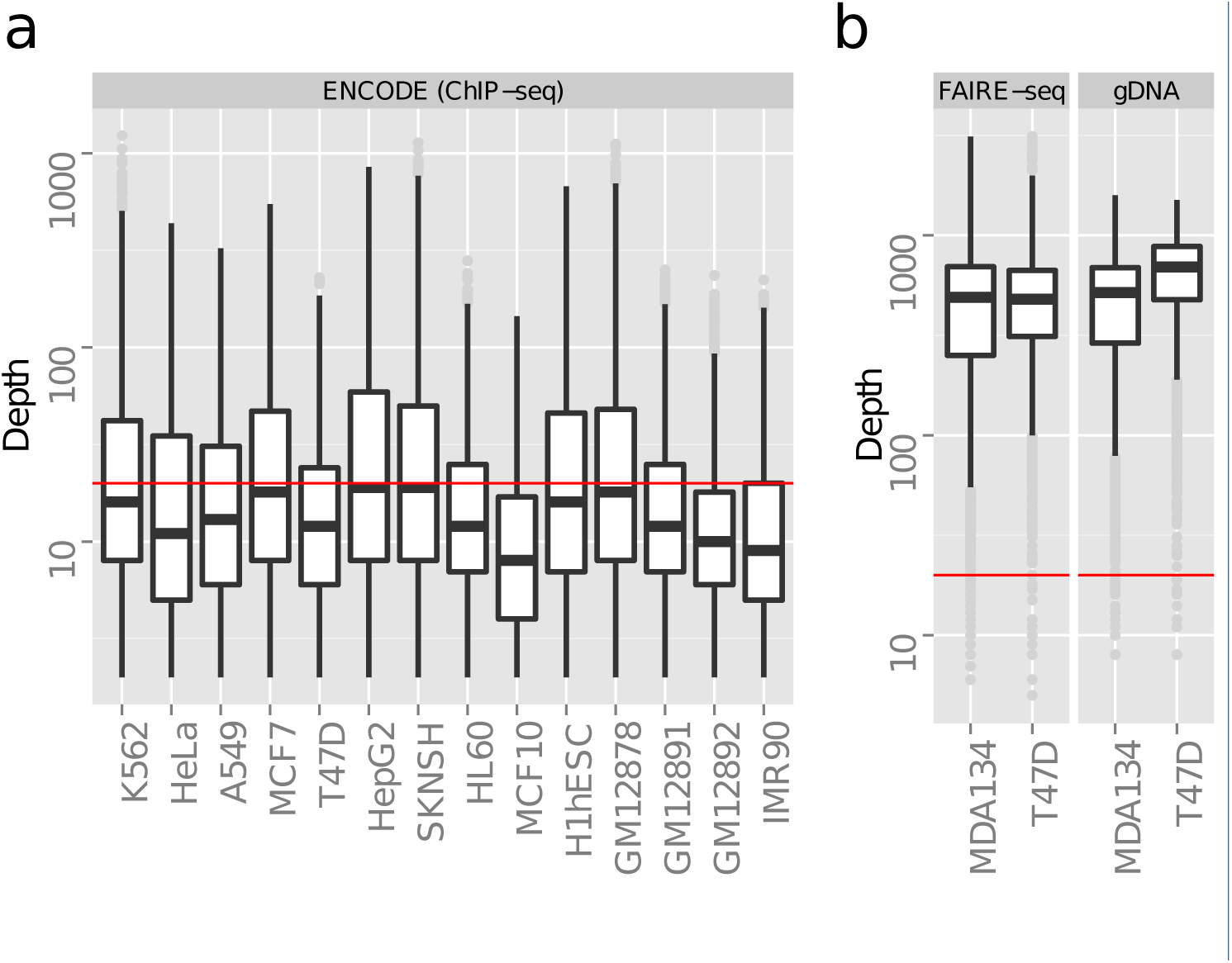
Distribution of depth of coverage at tested heterozygous SNPs in (a) ENCODE ChIP-seq datasets (pooled data across all ChIP-seq samples in each cell line) and (b) targeted sequencing FAIRE-seq datasets (pooled data for three replicates).

## Additional Files

Additional file 1 — supplementary_tables.xlsx

Supplementary tables S1 — S8.

